# Spatiotemporal transcriptomic atlas of mouse organogenesis using DNA nanoball patterned arrays

**DOI:** 10.1101/2021.01.17.427004

**Authors:** Ao Chen, Sha Liao, Mengnan Cheng, Kailong Ma, Liang Wu, Yiwei Lai, Xiaojie Qiu, Jin Yang, Wenjiao Li, Jiangshan Xu, Shijie Hao, Xin Wang, Huifang Lu, Xi Chen, Xing Liu, Xin Huang, Feng Lin, Zhao Li, Yan Hong, Defeng Fu, Yujia Jiang, Jian Peng, Shuai Liu, Mengzhe Shen, Chuanyu Liu, Quanshui Li, Yue Yuan, Huiwen Zheng, Zhifeng Wang, Zhaohui Wang, Xin Huang, Haitao Xiang, Lei Han, Baoming Qin, Pengcheng Guo, Pura Muñoz- Cánoves, Jean Paul Thiery, Qingfeng Wu, Fuxiang Zhao, Mei Li, Haoyan Kuang, Junhou Hui, Ou Wang, Haorong Lu, Bo Wang, Shiping Liu, Ming Ni, Wenwei Zhang, Feng Mu, Ye Yin, Huanming Yang, Michael Lisby, Richard J. Cornall, Jan Mulder, Mathias Uhlen, Miguel A. Esteban, Yuxiang Li, Longqi Liu, Xun Xu, Jian Wang

## Abstract

Spatially resolved transcriptomic technologies are promising tools to study cell fate decisions in a physical microenvironment, which is fundamental for enhancing our knowledge of mammalian development. However, the imbalance between resolution, transcript capture and field of view of current methodologies precludes their systematic application to analyze relatively large and three-dimensional mid- and late-gestation mammalian embryos. Here, we combined DNA nanoball (DNB) patterned arrays and tissue RNA capture to create SpaTial Enhanced REsolution Omics-sequencing (Stereo-seq). This approach allows transcriptomic profiling of large histological sections with high resolution and sensitivity. We have applied Stereo-seq to study the kinetics and directionality of transcriptional variation in a time course of mouse organogenesis. We used this information to gain insight into the molecular basis of regional specification, neuronal migration and differentiation in the developing brain. Furthermore, we mapped the expression of a panel of developmental disease-related loci on our global transcriptomic maps to define the spatiotemporal windows of tissue vulnerability. Our panoramic atlas constitutes an essential resource to investigate longstanding questions concerning normal and abnormal mammalian development.

## INTRODUCTION

Understanding how a single totipotent cell, the zygote, develops into a complex organism such as a mammal in a precisely controlled manner, over time and space, is one of the most fascinating questions in science ever since Aristotle. Mouse is an excellent model mammal to study developmental processes and establish parallels with larger species thanks to the inbred genetic background, susceptibility to genetic manipulation and relatively short gestation period (∼21 days). Recent intense efforts on cell atlas using high-throughput single-cell RNA-sequencing (scRNA-seq) technologies have provided an increasingly detailed view of mouse developmental gene expression dynamics (Cao et al., 2019; He et al., 2020). However, except for the early embryonic time points where fewer cells are present (Peng et al., 2019; Xue et al., 2013), topographical transcriptomic information is still mostly lacking. This deficiency complicates the interpretations, in particular the correlation between signaling cues and cell fate decisions. In this regard, embryonic decisions are often regulated by gene expression thresholds rather than binary mechanisms. The outstanding need for precise coordination of complex signals and the myriad embryonic cell types in development is exemplified by genetic disorders like situs inversus, where alterations in the response to nodal flow cause inversion of the internal anatomy (Okada et al., 2005). Systematic spatially resolved analysis of gene expression is thus strictly necessary to dissect the intricacies of mammalian development.

Several remarkable methodologies have been recently developed that allow spatially resolved transcriptomic profiling of tissue sections. They can be divided in three main classes based on their detection method: physical segmentation using microdissection coupled to sequencing (Nichterwitz et al., 2016; Peng et al., 2019), pre-designed probes for either single-molecule hybridization (Eng et al., 2019; Zhuang, 2021) or amplification and *in situ* decoding (Alon et al., 2021; Lee et al., 2014; Wang et al., 2018), and barcoded probes (Cho et al., 2021a; Rodriques et al., 2019; Srivatsan et al., 2021; Stahl et al., 2016; Vickovic et al., 2019; Yao et al., 2020). These techniques all have specific advantages and disadvantages in resolution and applicability but share a major caveat in the limited field of view. Although small parts of developing mouse tissues (Di Bella et al., 2021; Stickels et al., 2021) or even whole mouse embryos (Peng et al., 2019; Srivatsan et al., 2021; Yao et al., 2020) have been analyzed using these approaches, no available technology can profile samples of the size of whole mouse embryos in mid- or late-stage gestation with high definition. This poses a tremendous challenge for systematically studying the differentiation continuum that happens across different areas of the same tissue and for comparing different tissues simultaneously. Here, we report the development of a new DNB-based technology, Stereo-seq, that combines high resolution and sensitivity with large field of view to generate the first panoramic transcriptomic atlas of mouse organogenesis.

## RESULTS

### DNB patterned arrays enable large field-of-view spatially resolved transcriptomics with high definition

DNB sequencing is based on rolling circle amplification of DNA fragments (Drmanac et al., 2010), whose products are effectively docked into defined positions of a lithographically etched chip for *in situ* sequencing through automated high-throughput microscopic imaging. We envisaged that these features could lay the foundation for a spatially resolved transcriptomic technology with high-resolution and large field of view, which we developed as following. Standard DNB chips have spots with approximately 220 nm diameter and a center-to-center distance of 500 or 715 nm (Figure 1A**, step 1**). This provides up to 400 spots for tissue RNA capture per 100 µm^2^ (average size of a mammalian cell). DNB templates containing random barcodes are deposited on the patterned array, incubated with primers, and sequenced to obtain the data matrix containing the coordinate identity (CID) of every DNB (Figure 1A**, step 2**). The use of random barcode-labeled DNB allows a large spatial barcode pool size (4^25^). Next, unique molecular identifiers (UMI) and polyT sequence-containing oligonucleotides are ligated onto each spot through hybridization with an oligonucleotide sequence containing the CID (Figure 1A**, step 3**). Frozen tissue sections (10 µm thickness) are loaded onto the chip surface, followed by fixation, permeabilization to capture the tissue polyA-tailed RNA and finally reverse transcription plus amplification (Figure 1A**, step 4**). Amplified barcoded cDNA is collected, used as template for library preparation and sequenced together with the CID (Figure 1A**, step 5**). Computational analysis of sequencing data generated this way allows high-resolution spatially resolved transcriptomics (Figure 1A**, step 6**). We named this approach Stereo-seq (SpaTial Enhanced REsolution Omics-sequencing). Importantly, Stereo-seq has a larger number of spots per 100 µm^2^, with smaller spot size and center-to-center distance, than other available methods (Figure 1B). This makes it particularly suitable for detecting transcriptomic gradients with high resolution and sensitivity. So far, we have used Stereo-seq chips of up to an effective area of 13.2 cm x 13.2 cm for different tissue sizes, subdivided into 50 mm^2^, 100 mm^2^ and 200 mm^2^ smaller chips for profiling sections from the mouse olfactory bulb (∼10.5 mm^2^), a whole mouse brain (∼52.8 mm^2^) and whole mouse embryos (∼7.1 to ∼97.6 mm^2^), respectively (Figure S1A). These capture areas are substantially larger than those achieved by other available technologies (Figure 1B), and we anticipate the application of unsliced chips to much larger tissues such as whole human brain sections (Figure S1A).

**Figure 1.**
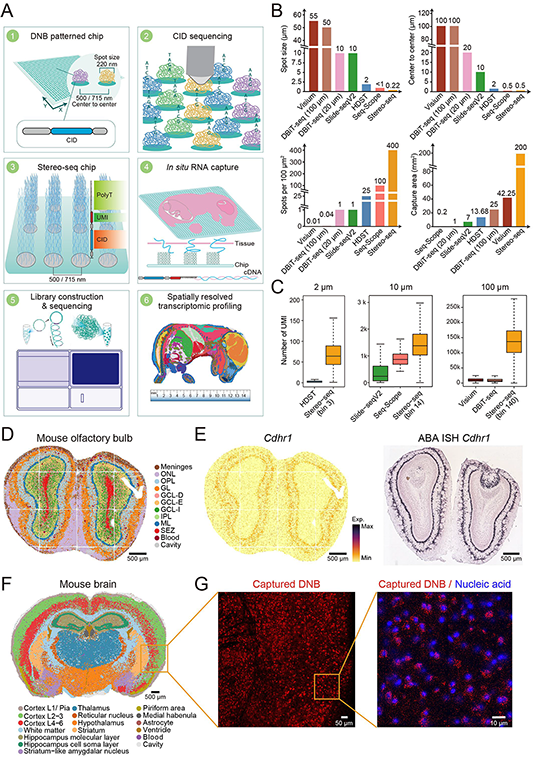
Advanced RNA capture from complex tissues by Stereo-seq. **A.** Stereo-seq pipeline. *Step 1*, design of the DNB patterned array chip. *Step 2*, *in situ* sequencing to determine the spatial coordinates of uniquely barcoded oligonucleotides placed on each spot of the chip. *Step 3*, preparation of capture probes by ligating the UMI-polyT containing oligonucleotides to each spot. *Step 4*, subsequent *in situ* RNA capture from tissue placed on the chip. *Step 5*, cDNA amplification, library construction and sequencing. *Step 6*, data analysis to generate the spatially resolved transcriptome of the profiled tissue. **B.** Stereo-seq achieves a smaller spot size (upper left), higher resolution (upper right), higher number of spots per 100 μm^2^ (bottom left) and larger capture area (bottom right) than other reported methods. Samples used for the comparison included mouse olfactory bulb (Stereo-seq, Visium, Slide-seqV2 and HDST), E10 mouse embryo (DBiT-seq) and mouse liver (Seq-Scope) (Cho et al., 2021b; Lebrigand et al., 2020a; Liu et al., 2020; Stickels et al., 2021; Vickovic et al., 2019). Note that since Seq-Scope uses a random array, which contains no patterned spots, the size of each pixel was estimated according to the published dataset. **C.** Boxplots showing the number of transcripts captured by Stereo-seq at the indicated resolution in comparison with reported HDST, Slide-seqV2, Visium, DBiT-seq and Seq-Scope datasets. Samples in those datasets used for comparison are as in **panel B**. **D.** Unsupervised spatially-constrained clustering of the mouse olfactory bulb section analyzed by Stereo-seq data at bin 14 resolution, bins were colored by their annotation. ONL, olfactory nerve layer. OPL, outer plexiform layer. GL, glomerular layer. GCL-D, granular cell zone deep. GCL-E, granular cell layer externa. GCL-I, granular cell layer internal. IPL, internal plexiform layer. ML, mitral layer. SEZ, subependymal zone. Scale bar, 500 μm. **E.** Left: spatial visualization of *Cdhr1* in a mouse olfactory bulb section analyzed by Stereo-seq. Right: *Cdhr1* ISH data taken from Allen Brain Atlas. Scale bars, 500 μm. **F.** Unsupervised spatially-constrained clustering of the mouse brain section analyzed by Stereo-seq at bin 50 resolution, bins were colored by their annotation. Scale bar, 500 μm. **G.** Left: projection of captured DNB signals of the same region squared in the **panel F**. Scale bar, 50 μm. Right: superimposed nucleic acid staining and captured DNB signals from the same region squared in the middle panel. Scale bar, 10 μm.

To benchmark our technology, we first profiled the mouse olfactory bulb because it is a widely used model tissue for spatially resolved transcriptomic approaches (Lebrigand et al., 2020b; Rodriques et al., 2019; Vickovic et al., 2019). Stereo-seq captured numbers ranging on average 69 UMI per 2 µm (diameter) bin (bin 3, 3 × 3 DNB), 1,450 UMI per 10 µm (diameter) bin (bin 14, 14 × 14 DNB, equivalent to ∼1 medium size cell) and 133,776 UMI per 100 µm (diameter) bin (bin 140, 140 × 140 DNB) (Figure 1C **and** S1B). This is superior to other available technologies including Seq-Scope (848 UMI on average per 10 µm diameter bin) (Cho et al., 2021a) when compared at the same bin resolution.

Conventional cell taxonomy analysis with single-cell RNA-seq (scRNA-seq) uses algorithms such as Leiden clustering to group cells based on transcriptome similarity (Traag et al., 2019). Applying this method to spatially resolved data, however, is problematic due to the lack of consideration of spatial coordinates. Thus, we developed a spatially-constrained clustering algorithm especially suited for high-resolution spatially resolved transcriptomic data (**see Methods**), which clustered bins over a continuous area from which tissue domains are annotated. This approach showed high level of consistency between biological replicates corresponding to two adjacent sections of the mouse olfactory bulb (R^2^ = 0.963) (Figure 1D, S1C, **and** S1D). The distribution of specific markers (*Cdhr1*, *Pcp4*, *Slc17a7*, *Sox11* and *Th*) captured by Stereo-seq exhibited remarkable similarity to *in situ* hybridization (ISH) data (Lein et al., 2007), and showed clearer patterns than HDST or Slide-seqV2 (Figure 1E, S1C, **and** S1E).

To further demonstrate the robustness of Stereo-seq and prove its scalability, we profiled an entire adult mouse coronal brain section. Unsupervised spatially-constrained clustering of the Stereo-seq data identified different anatomical structures including the cortex and subcortical regions such as hippocampus, thalamus and striatum (Figure 1F). This is so far the largest brain spatial atlas profiled by a genome-wide sequencing based spatial technology. Differentially expressed gene (DEG) and gene ontology (GO) enrichment analyses confirmed the unique identity of the observed structures (Figure S2A and S2B). When focusing on specific elements such as the hippocampus (*Ncdn^+^* and *Neurod6^+^*), thalamus (*Prkcd^+^*), hypothalamus (*Sparc^+^*) and striatum (*Penk^+^*), among others, we observed specific spatial distribution of known markers (Figure S2C). Notably, we used a specific dye for nucleic acids before *in situ* permeabilization to assess the resolution of the data after sequencing. When zooming into a specific region corresponding to the cortex, we observed substantial colocalization of the dye and the aggregated transcripts detected by Stereo-seq (Figure 1G). This confirmed that Stereo-seq captures transcripts at cellular resolution.

We concluded that Stereo-seq can be used to spatially characterize the transcriptomic pattern of complex tissues in an unbiased multiplexed manner with high definition and unprecedented field of view.

### Spatially resolved transcriptomic atlas of mouse organogenesis using Stereo-seq

We postulated that the unique features of Stereo-seq would allow us to create panoramic transcriptomic maps of middle- and late-gestation mouse embryos for which full body sections cannot be profiled using other available technologies. This would also showcase the strength of Stereo-seq in measuring spatiotemporal transcriptional dynamics at full scale. To this aim, we created the beginning of Mouse Organogenesis Spatiotemporal Transcriptomic Atlas (MOSTA), an expandable and interactive database for spatial transcriptomic studies of mouse organogenesis. As proof of principle, we profiled 14 sagittal sections (two sections per embryo) from C57BL/6 mouse embryos at E9.5 (∼7.1 mm^2^), E10.5 (∼10.9 mm^2^), E11.5 (∼19.7 mm^2^), E12.5 (∼30.2 mm^2^), E13.5 (∼55.2 mm^2^), E15.5 (∼93.2 mm^2^) and E16.5 (∼97.6 mm^2^). These stages cover most of the key events in mouse organogenesis (Cao et al., 2019).

To gain an unbiassed global view of MOSTA, we segmented our individual embryo datasets into 50 × 50 DNB bins (bin 50 resolution, 36 µm diameter). In total, we retrieved transcriptomic information for 616,125 bins. The average number of captured genes per bin in each embryo ranged from 1,088 at E12.5 to 3,153 at E11.5, and of UMI from 3,473 at E13.5 to 10,939 at E11.5 (Figure S3A). Unsupervised spatially-constrained clustering showed the expected major organs and systems (e.g., nervous system, heart, liver, lung, kidney, adrenal gland, gastrointestinal tract, genitourinary tract, skin, bone and muscle) at each respective time point, whose external boundaries closely resembled the anatomical structures (Figure 2A, S3B, **and** S3C). DEG analysis showed the expected correspondence with emerging tissue-specific cell identities at individual stages (Figure S4). This was further confirmed by plotting specific marker genes on the spatial maps across all embryonic time points (Figure 2B **and** S5). Since Stereo-seq captures very high density of signals, we also performed spatial re-clustering of bins for selected tissues, which produced detailed structures containing different subregions (Figure 2C). For example, re-clustering of the E10.5 heart identified the atrium (*Myh6*^+^) and ventricle (*Myh7*^+^/*Myl2*^+^); of the E15.5 limb the cartilage primordium (*Col2a1*^+^), dermis (*Dcn*^+^) and epidermis (*Krt15*^+^); of the E16.5 spinal cord the marginal (*Scng*^+^), mantle (*Snrpn*^+^/*Plxna2*^-^) and ventricular zones (*Snrpn*^+^/*Plxna2*^+^); and of the E16.5 adrenal gland the medulla (*Dbh*^+^), cortex (*Fdx1*^+^) and capsule (*Col1a2*^+^). Notably, reanalysis of E10 mouse embryo DBiT-seq data (Liu et al., 2020) showed significantly fewer 50 µm-diameter spots (902) than 36 µm diameter (bin 50)-bins (7,386) were calculated using Stereo-seq in an individual E10.5 section. Consequently, unsupervised clustering or mapping of specific genes failed to identify discernable anatomical structures with DBiT-seq (Figure S6A and S6B). This discrepancy is related to the large inter-channel space (50 µm) of DBiT-seq.

**Figure 2.**
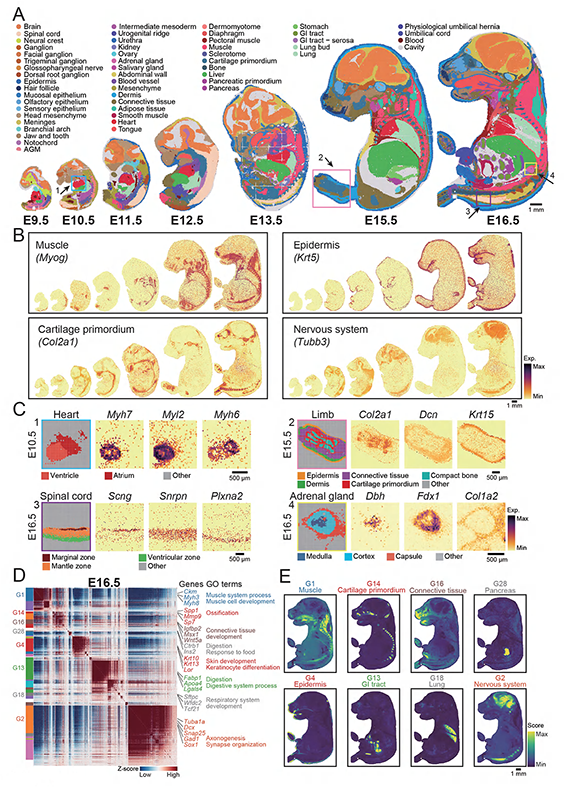
Spatiotemporal transcriptomic atlas of mouse organogenesis. **A.** Unsupervised spatially-constrained clustering of mouse embryo sections across E9.5, E10.5, E11.5, E12.5, E13.5, E15.5, and E16.5, bins were colored by their annotation. The squares indicate the regions for analysis in **panel C**. Scale bar, 1 mm. **B.** Spatial visualization of expression of indicated genes for muscle, epidermis, cartilage primordium, and nervous system. Scale bar, 1 mm. **C.** Spatial visualization of indicated areas in **panel A** identified detailed anatomical structure for the heart (E10.5), limb (E15.5), spinal cord (E16.5), and adrenal gland (E16.5). The expression of marker genes for each anatomical structure is shown on the right side of each area. Scale bars, 500 μm. **D.** Heatmap showing the genes with significant spatial autocorrelation (3,240 genes, FDR < 0.05) grouped into 28 gene modules on the basis of pairwise spatial correlations of E16.5 embryo. Selected genes and GO terms related to representative gene modules are highlighted on the right side. **E.** Spatial visualization of indicated gene module related to muscle, cartilage primordium, connective tissue, pancreas, epidermis, GI tract, and nervous system. Scale bar, 1 mm.

To study cell function enrichments within specific areas contained in the individual tissue clusters generated with Stereo-seq, we applied Hotspot (DeTomaso and Yosef, 2021), an algorithm that measures non-random variation to recognize informative gene programs. The latter is important because amalgamation of signals from different cell types and states in complex tissues such as the embryo can otherwise complicate the interpretations. We identified 28 gene modules for E9.5, 28 for E10.5, 33 for E11.5, 30 for E12.5, 24 for E13.5, 23 for E15.5 and 28 for E16.5 (Figure 2D, 2E**, and** S7-S13; **Table S1**). GO enrichment analysis demonstrated that most of these spatial gene modules correspond to features related to region-specific biological processes such as muscle development (module G1), neurogenesis (module G2), skin development (module G4), digestive system (module G13) and ossification (module G14) at E16.5 (Figure 2D, 2E**, and** S13A; **Table S1**). Then, we used a set of manually annotated cell types from a recent scRNA-seq atlas of E9.5-E13.5 mouse embryos (Cao et al., 2019) to assign putative cell identities to our spatial modules, observing good correlation. For example, genes specific to hepatocytes and erythroid lineage, megakaryocytes or neutrophils showed strong co-enrichment in the same module starting from E11.5 but were not earlier, consistent with the fact that the definitive hematopoiesis only occurs in the liver from E11.5 onwards (Lewis et al., 2021) (Figure S9C). Likewise, radial glial cells are the source of neuroblasts (Merkle et al., 2004) and both cell types co-localized in brain regions since E11.5. In addition, jaw and tooth progenitor cells co-localized with osteoblasts and connective tissue progenitor cells also after E11.5. The complete set of cell assignments for all tested time points can be found in Figure S7-S13.

Therefore, MOSTA generated by Stereo-seq constitutes a unique resource to investigate the molecular basis of tissue patterning and the emergence of cell identities during mouse organogenesis. Our publicly available interactive data portal can be accessed at https://db.cngb.org/stomics/mosta/.

### Spatiotemporal dynamics of gene regulation in mouse organogenesis

Cell identity maintenance and cell state dynamics are governed by intrinsic and inducible gene regulatory networks (Aibar et al., 2017). These networks are often studied in the form of modules of co-expressed genes that are co-regulated by common sets of transcription factors, or regulons (Aibar et al., 2017). Although this data can be extracted from scRNA-seq datasets applying algorithms such as SCENIC (Aibar et al., 2017), the lack of spatial information muddles the establishment of reliable associations. We used SCENIC to calculate the frequencies of predicted transcription factor binding events in each of the individual time points of MOSTA. This identified many regulons presenting regional specificity at each stage. To achieve a global overview of these transcriptional programs and how they are interconnected, we applied again Hotspot, which categorized them into a series of regulon modules (**Table S2**). Many of these modules contained sets of transcription factors known to be involved in the formation of specific tissues and/or specific developmental processes (Figure 3A **and** S14-S19). For example, at E9.5 we identified 23 modules containing a total of 410 regulons corresponding to different tissues (Figure 3A). Among these, GATA4 was enriched in the heart (module R1), GATA1 in the blood (module R2), SOX2 in the brain (module R4), DLX6 in the branchial arch (module R10) FOXA2 in the notochord (module R11) and HNF4A in the liver (module R18) (Figure 3B).

**Figure 3.**
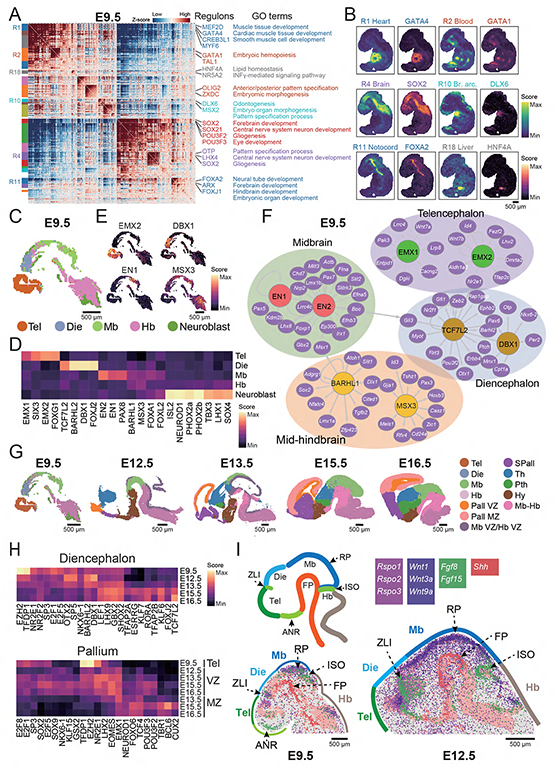
Spatially resolved gene regulatory networks of organogenesis. **A.** Heatmap showing the regulons with significant spatial autocorrelation (481 regulons, FDR < 0.05) grouped into 23 modules on the basis of pairwise spatial correlations of E9.5 embryo section, selected regulons and their corresponding GO terms related to representative regulon modules are highlighted on the right side. **B.** Spatial visualization of indicated modules and their representative regulon for heart, blood, brain, branchial arch, notochord, and liver. Scale bar, 500 μm. **C.** Unsupervised spatially-constrained clustering of E9.5 mouse embryonic brain based on regulon activity. Tel: telencephalon; Die: diencephalon; Mb: midbrain; Hb: hindbrain. Scale bar, 500 μm. **D.** Heatmap showing the normalized regulon activity score of top transcription factors corresponding to different anatomical structures of E9.5 brain. **E.** Spatial visualization of the indicated regulon activity representing different anatomical structures of E9.5 brain. Scale bar, 500 μm. **F.** Gene regulatory network of transcription factors (EMX1/2) representing telencephalon (EMX1/2), diencephalon (TCF7L2 and DBX1), midbrain (EN1/2), and mid-hindbrain (BARHL1 and MSX3) of E9.5 brain as visualized by Cytoscape. Selected target genes related to neuronal development was shown. **G.** Unsupervised spatially-constrained clustering of mouse embryonic brain from E9.5, E12.5, E13.5, E15.5 and E16.5 based on regulon activity identified anatomical structures of the developing brain. MZ: mantle zone; VZ: ventricular zone; Pall, pallium; Spall, subpallium; Th, thalamus, Pth, prethalamus; Hy, hypothalamus; Mb-Hb, mammillary body - hindbrain. Scale bars, 500 μm. **H.** Heatmaps showing the normalized regulon activity score of top transcription factors corresponding to mouse embryonic diencephalon (top) or pallium (bottom) development. **I.** Upper left: schematic representation of the secondary organizer centers in the early embryo brain. Bottom: Spatial scatter plots showing single DNB that captured the transcripts for the indicated morphogens at the ISO, ANR, ZLI, RP, and FP at E9.5 (bottom left) and E12.5 (bottom right) embryos. Plots were superimposed with E9.5 or E12.5 brain structures. ISO, isthmus organizer; ANR, anterior neural ridge; ZLI, zona limitans intrathalamica; RP, roof plate; FP, floor plate. Scale bars, 500 μm.

We focused on the brain for more detailed investigation because the relatively large dimensions, unparalleled complexity and prolonged developmental time makes particularly challenging the study of regulons with single-cell sequencing approaches. First, we performed unsupervised clustering of E9.5 brain regulons. This showed four major neuroepithelial clusters (*Pax6^+^* and/or *Sox2^+^*) corresponding to the telencephalon, diencephalon, midbrain, and hindbrain, with a smaller neuroblast-like cluster (*Nhlh1/2*^+^) distributed in the diencephalon and more obviously in the hindbrain (Figure 3C **and** S20A). Several transcription factors with known functions in brain development showed differential activity among these regions (Figure 3D-3F **and** S20B-S20D). For example, the telencephalon was enriched in EMX1/2, which regulate proliferation and specification of radial glial cells (Bishop et al., 2003). The diencephalon was enriched in transcription factors involved in thalamus specification (e.g., TCF7L2, BARHL2 and DBX1) (Lipiec et al., 2020; Vue et al., 2007). EN1/2 were enriched in the anterior and posterior midbrain and are known to regulate dopaminergic neuron specification (Alves dos Santos and Smidt, 2011). Mid- and hindbrain shared some regulons including BARHL1, which is known to be specifically enriched in the adult cerebellum (part of the hindbrain) (Li et al., 2004), and LMX3. Likewise, the early neuroblast regulon was enriched in transcription factors characteristic of this cell type (e.g., SOX4, NEUROD1 and LHX1) (Liu et al., 2000) (Figure 3D **and** S20B).

We next systematically examined the regulons that direct regional brain specification across different time points. We focused on E9.5 and E12.5-E16.5 because of the enhanced capture of related structures. We identified regulon clusters controlling the formation and expansion of the diencephalon, pallium, subpallium, midbrain, and hindbrain (Figure 3G). In the diencephalon, the neuroepithelial regulons (e.g., DBX1 and BARHL2) identified at E9.5 became progressively inactivated after E12.5, whereas TCF7L2, BARHL2 and DBX1 increase until E16.5 (Figure 3H). Another group of regulons comprising factors associated with thalamus maturation (e.g., RORA and TFAP2B) (Vitalis et al., 2018) appeared at E15.5 and remained high at E16.5. Several regulons were shared between all stages (e.g., LEF1 and SHOX2) but their activity increased towards E16.5, suggesting that fine-tuning their levels orchestrates the transition from specification to maturation. Regulons in the pallium (originated from the telencephalon) at later stages were divided into the ventricular zone (VZ) and mantle zone (MZ) (Figure 3G). E9.5 telencephalon regulons progressively declined after this time point (Figure 3H). Many of the VZ-specific regulons showed transcription factors related to early neurogenesis (e.g., EOMES, LHX2 and E2F family members) (Greig et al., 2013) and remained high from E12.5 onwards (Figure 3H). In contrast, those involved in MZ formation were related to neuronal maturation (e.g., CUX2 and BCL6), appeared at E13.5 and increased until E15.5 or E16.5. Subpallium regulons showed strong enrichment of DLX homeobox transcription factors (DLX1/2/5/6) from E13.5 to E16.5 (Figure S20E **and** S20F), consistent with their role in GABAergic interneuron production (Sultan et al., 2013). Midbrain and hindbrain largely showed similar sets of regulons associated with neuron progenitor at early stage (e.g., E2F1, E2F5 and SP3), but presented different set of transcription factors related to neuronal maturation at later stages in each region (e.g., LEF1 and DMBX1 in the midbrain, and HOXA5 and HOXB5 in the hindbrain) (Figure S20E and S20F). We next investigated the spatial expression of diffusible extracellular cues (morphogens) at the so-called secondary organizer centers of the developing brain. These morphogens are responsible for inducing changes in gene regulatory programs that cause regionalization in the brain (Tannahill et al., 2005). We studied the earlier stages (E9.5 and E12.5) because the biggest morphogenetic events of the brain happen during that period. Analysis of gene modules identified with Hotspot showed good correspondence with the known location of key secondary organizer centers including the floor plate (FP), roof plate (RP), anterior neural ridge (ANR), zona limitans intrathalamica (ZLI) and isthmus organizer (ISO) (Figure S21A **and** S21B). Spatial correlation analysis of ligands in these gene modules revealed distinct clusters in each stage (Figure S21C **and** S21D). These morphogen clusters included ligands from the Sonic hedgehog (SHH), Wnt, bone morphogenetic proteins (BMP), fibroblast growth factors (FGF) and R-spondin (RSPO) families (Tannahill et al., 2005). We used this information to generate a spatial morphogen map of the developing brain (Figure 3I **and** S21E). Examples of morphogens involved in dorsoventral patterning included *Wnt1*, *Wnt3a*, *Wnt9a* and *Rspo1/2/3* at the RP, whereas *Shh* was mainly enriched at the FP and to a less extent at the ZLI. Morphogens involved in anteroposterior patterning included FGF family members such as *Fgf8* and *Fgf15* at the ANR, ZLI and ISO. We noticed that regulons at these secondary organizers were quite different and variable during development. For example, EN1/2 were enriched at the ISO in both E9.5 and E12.5 embryos, while the ZLI was enriched in DMBX1 and PAX6 at E9.5 or PRDM16 and MAFF at E12.5. In addition, FOXA1/2/3 were enriched at the FP and OTX2 in the RP (Figure S22). These results highlight the utility of Stereo-seq to study coordinated transcriptional responses to signaling cues at large scale.

Regulons for other tissues also showed a gradual spatiotemporal transition. For example, in the skeleton we identified regulons consisting of the early condensation regulators SOX5 and SOX9 (Kozhemyakina et al., 2015), which promote the transition from mesenchymal progenitors in the sclerotome (E9.5-E11.5) to the early (cartilage primordium, E12.5 and E13.5) and late (chondrocyte, E15.5 and E16.5) chondrogenic linage (Figure S23A-S23C). Likewise, we detected key regulators of chondrocyte to osteoblast maturation such as SP7 and XBP1 (Kozhemyakina et al., 2015), which correlated with a switch in the pattern of extracellular matrix production from *Col9a3* to *Col22a1*.

In summary, MOSTA can be used to infer the gene regulatory networks behind patterning events driving organogenesis. Systematic analysis of other regulons, morphogens and tissues can be performed using our interactive portal.

### Regulation of cell diversification in the telencephalon

The precise regulation of neuronal cell fate is critical for the development of brain architecture, of which the neocortex is particularly interesting because of the diversity of neuronal subtypes and the sophisticated migration trajectories (Greig et al., 2013). We first focused on the late stage telencephalon (E15.5), when the pallium and subpallium become apparent. Unsupervised spatially-constrained clustering at bin 20 resolution (20 × 20 DNB, 14 μm diameter) identified clusters corresponding to known anatomical regions (Figure 4A). These clusters displayed specific markers (Figure S24A **and** S24B). Briefly, we could clearly distinguish distinct pallium clusters including the dorsal pallium (DPall), medial pallium (MPall, also known as hippocampal allocortex) and ventral pallium (VPall). The DPall included clusters corresponding to the VZ, subventricular zone (SVZ), intermediate zone (IZ), cortical plate (CP) and marginal zone (MarZ). All subpallium regions expressed markers related to the generation and maturation of cortical interneurons (e.g., *Dlx5* and *Gad2*; Figure S24A **and** S24B) (Greig et al., 2013), and also markers corresponding to known anatomical areas including the central subpallium (CSPall) and paraseptal subpallium (PaSe) (https://developingmouse.brain-map.org/static/atlas). Through enrichment of known genes, we further located the striatum and lateral ganglionic eminence (LGE) (*Bcl11b, Foxp1/2, Isl1* and *Ebf1*), medial ganglionic eminence (MGE) and preoptic area (POA) (*Nkx2.1, Lhx6* and *Lhx8*) (Figure 4A **and** S24C). Of note, the spatial clustering showed two clusters along the rostral to caudal direction at the CP (Figure 4A). DEG analysis revealed graded distribution of genes related to axonogenesis and neuron projection development along this axis (Figure S25A **and** S25B), reflecting different kinetics of projection neuron specification from partially fate-restricted postmitotic progenitors (Greig et al., 2013). Interestingly, we also observed a gradient of expression for morphogens such as *Nrg3* along the rostral to caudal direction (Muller et al., 2018; Nowakowski et al., 2017) (Figure S25C **and** S25D**; Table S3A)**.

**Figure 4.**
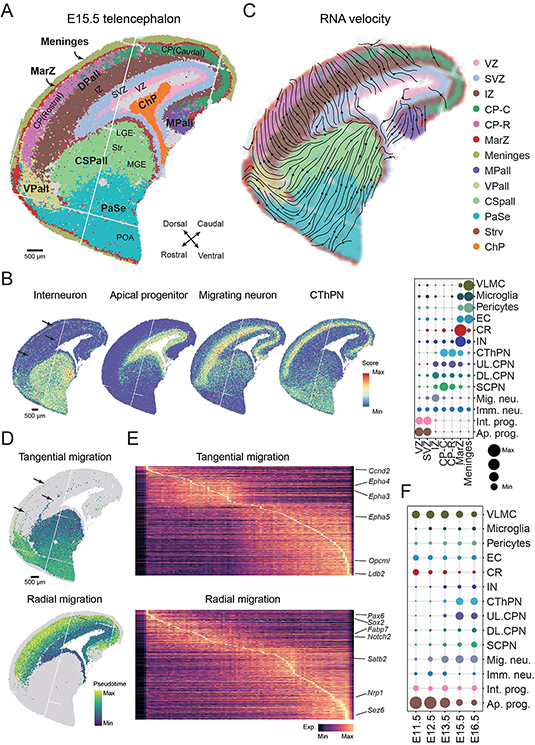
Cell type diversification during neocortex development. **A.** Unsupervised spatially-constrained clustering of the forebrain from E15.5 embryo identified anatomical structures (bin 20 resolution). Bins are colored by their annotated structures. CP-C, cortical plate - caudal; CP-R, cortical plate - rostral; MarZ, marginal zone; MPall, medial pallium; VPall, ventral pallium; CSPall, central subpallium; PaSe, paraseptal subpallium; StrV, ventricular zone of subpallium; ChP, choroid plexus. Scale bar, 500 μm. **B.** Left: spatial visualization of the indicated cell type proportion score for the E15.5 forebrain. Right: dotplot showing the cell type composition at different anatomical structures of the E15.5 forebrain. Ap. Prog, apical progenitor; Int. prog., intermediate progenitor; Imm. neu., immature neurons; Mig. Neu. migrating neuron; SCPN, subcerebral projection neurons; DL. CPN, deeper layer callosal projection neuron; UL. CPN, upper layer callosal projection neuron; CThPN, corticothalamic projection neuron; IN, Interneuron. CR, Cajal-Retzius cell. EC, endothelial cell. VLMC, vascular and leptomeningeal cell. Scale bar, 500 μm. **C.** RNA velocity streamline plots visualize the developmental trajectory of the forebrain. Bins are colored by their annotated structures, as in **panel A**. Scale bar, 500 μm. **D.** Vector field pseudotime analysis of tangential migration of interneuron neurons (upper) and radial migration of cortical neuron (bottom). Scale bar, 500 μm. **E.** Gene expression heatmap of 680 top PCA genes in a pseudo-temporal order of tangential migration (upper) and radial migration path (bottom), genes with highest PCA loading as well as the DEG identified by unsupervised spatially-constrained clustering were shown in the figure. **F.** Dotplot showing the cell type composition dynamic for the mouse embryonic cortex at E11.5, E12.5, E13.5, 15.5, and E16.5.

We then applied SPOTlight (Elosua-Bayes et al., 2021) to map the cell types into the spatial maps using a recently published neocortex single-cell dataset (Di Bella et al., 2021). This detected a variety of neuronal and non-neuronal cell types (Figure 4B **and** S26A). The VZ and SZV largely showed radial glial cells and intermediate progenitor cells, which decline at the IZ being replaced by immature neurons. For cortical projection neurons, we observed corticothalamic projection neuron (CThPN) in the inner CP in the vicinity of immature neurons, and subcerebral projection neurons (SCPN) mainly in the outer CP and more sparsely in the IZ. Consistent with our findings above, we observed higher level of upper layer cortical projection neurons (UL.CPN) in the rostral cluster of the CP, while deeper layer cortical projection neurons (DL.CPN) had similar distribution.

To detect the directionality of gene expression changes in the developing telencephalon, we applied Dynamo (**see Methods**), an analytical framework that measures transcriptional dynamics to predict the emergence of cell fates via vector field reconstruction (Qiu et al., 2021). In this regard, over 10% of the Stereo-seq captured transcripts contained intronic reads (Figure S26B), comparable to the proportion captured by scRNA-seq (La Manno et al., 2018). Spatial RNA velocity analysis revealed two strong directional flows of transcriptional changes, one from VZ and SVZ towards the CP, and the other from the subpallium to the cortical layers above it. The former direction is consistent with the path of neuronal specification from progenitor cells migrating radially towards the MZ, while the latter is consistent with the tangential migration of interneurons from the MGE into the neocortex at specific layers including the SVZ and IZ (Lodato and Arlotta, 2015) (Figure 4C). This agrees with the observed spatial probability score of cortical interneurons, projection neurons and neuron progenitors (Figure 4B). To study the regulators that guide interneuron and cortical neuron migration, we performed pseudotime analysis along the migration paths (Figure 4D and 4E**; see Methods)**. Interestingly, this showed both known regulators and unknown potential regulators of neuronal migration. For example, the ephrin receptor tyrosine kinases *Epha3, Epha4* and *Epha5*, were identified in the tangential path of interneurons (Bevins et al., 2011) (Figure S26C). Like wise, for radial migration, we identified *Sox2* and *Pax6* associated with early neurogenesis, and *Satb2* with projection neuron maturation (Alcamo et al., 2008; Gomez-Lopez et al., 2011). Other identified regulators are listed in **Table S3** and could be important for further study.

We next investigated the dynamic changes in projection neurons and interneurons across developmental stages in neocortex regions from E11.5 to E16.5 embryos. Unsupervised spatially-constrained clustering revealed clusters corresponding to specific layers in each group (Figure S27A). Single-cell projection into these spatial maps revealed striking cell type diversification during these stages. We observed that interneurons appear in the CP at E13.5 and expand at E15.5 and E16.5. In contrast, CPN appeared early (E12.5) and expanded and concentrated later at the IZ at E15.5 and E16.5. SCPN appeared at E12.5 and resided at the upper CP layer at later stages, while CThPN appeared later and resided at the inner CP layer (Figure 4F, S27B**, and** S27C). Cell cycle analysis further confirmed the dynamics from early progenitor to later postmitotic transition at the CP (Figure S27D).

Our results thus support the existence of spatiotemporal hierarchies governing cell-type diversification in the developing telencephalon (Nowakowski et al., 2017). Further scrutiny of this dataset using MOSTA database will be invaluable for defining associations with potential functional significance in the brain or other tissues.

### Deciphering developmental disease susceptibility using MOSTA

Developmental diseases have been classically regarded as those producing gross neonatal manifestations. Yet, many genetic conditions only manifested after birth have a developmental, often unnoticed, component because their target genes are also expressed during development (Boycott and Ardigo, 2018). For example, although Huntington’s disease is a late-manifesting neurological disorder affecting mostly the striatum, it is also associated with abnormalities in cortex development (Barnat et al., 2020). Nowadays, scRNA-seq datasets are becoming widely used to predict the origin and mechanisms of genetic diseases including developmental diseases (Watanabe et al., 2019). Yet, the cell capture bias and lack of contextual microenvironment affects the conclusions.

Given its unique characteristics, MOSTA could be particularly useful to dissect the developmental origins of mammalian genetic diseases, especially for understanding what tissue regions are originally affected and when, potentially shedding light into both big and subtle phenotypes. A major advantage compared to data generated with other spatially resolved technologies is the ability to look at multiple areas of the same tissue and multiple tissues simultaneously across different embryonic stages. To showcase this, we first took the 1,987 disease list (including 1,940 genes) from the Deciphering Developmental Disorders (DDD) database (Firth et al., 2011), filtered them to select the top 729 genes based on the expression threshold in our own dataset (**Table S4**), and projected both the filtered and unfiltered genes onto our spatial transcriptomic maps for all developmental time points. This showed enrichment in different tissues including the brain, heart, liver, blood, bone, muscle and others (Figure 5A **and** S28). Some of these disease-related genes were expressed in most or all embryonic stages, whereas others had a restricted window. Similarly, although many genes were mainly enriched in one tissue (in some cases confined to a specific region) or system, others were shared between several. For example, *Wnt5a*, which is related to Robinow syndrome, located in the craniofacial region, genital ridge, and brain (FP and neocortex) since early stages, consistent with the disease phenotypes (Menezes et al., 2010) (Figure 5B). *NKX2-1*, mutated in brain-lung-thyroid syndrome (Shetty et al., 2014), was mainly enriched in brain (hippocampus and subpallium) and lung. *Lmod3*, which is associated with nemaline myopathy (Berkenstadt et al., 2018), was enriched in the skeletal muscle.

**Figure 5.**
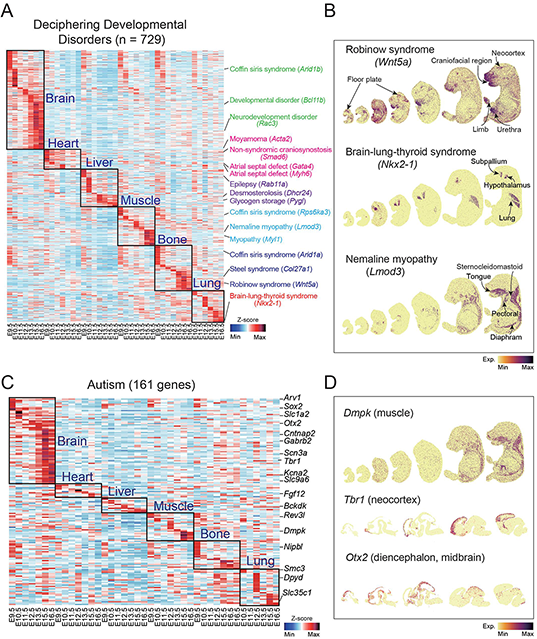
Association of MOSTA profiles with human developmental disorders. **A.** Heatmap showing the normalized expression level of 729 genes selected from the Deciphering Developmental Disorders database in six representative anatomical structures (brain, bone, muscle, heart, liver and lung) in embryos from E9.5 to E16.5. **B.** Spatial visualization of expression of indicated genes related to the human developmental disorders (Robinow syndrome, brain-lung-thyroid syndrome and nemaline myopathy) in mouse embryos from E9.5 to E16.5. Disease-related structures are annotated in the figure. **C.** Heatmap showing the normalized expression level of 161 autism-related genes in six representative anatomical structures (brain, bone, muscle, heart, liver and lung) in mouse embryos from E9.5 to E16.5. **D.** Spatial visualization of specific genes associated with autism (*Dmpk*, *Tbr1*, and *Otx2*) in the embryo brain from E9.5 to E16.5. Disease-related structures are annotated in the figures.

Autism encompasses a complex and multifactorial (genetic and environmental) group of neuropsychiatric diseases with recognized developmental roots but poorly understood mechanisms (Lord et al., 2020). Clarifying the diversity of tissues affected by autism besides the brain is important for designing patient-specific management therapies, as it is often associated with non-behavioral manifestations. We projected a battery of 161 autism-related genes (Liberzon et al., 2015) onto our spatial embryo maps, observing grouping in different tissues (Figure 5C; **Table S4**). Approximately 1/3 of the genes were enriched in the central nervous system (CNS), mostly the brain, with predominance or not for certain regions. Some autism-related genes were expressed along the whole developmental timeline, whereas others were more restricted (Figure 5C). For example, *Otx2* was enriched in the midbrain, subthalamus and pallium at different stages, whereas *Tbr1* was located mainly in the MZ of the pallium. Interestingly, a large group of autism related genes were enriched in non-neural tissues (e.g., heart, liver, muscle, bone and lung). For example, *Dmpk* was widely expressed in muscle but not in the brain, suggesting that abnormalities of the former tissue indirectly cause the autism phenotype (Figure 5D).

These results demonstrate the utility of MOSTA for defining the spatiotemporal windows of genetic disease vulnerability during development, and for establishing expected and unexpected causal relationships. Manual inclusion of other disease datasets can be used for case-specific explorations.

## DISCUSSION

Topographic transcriptomic information is necessary to dissect the molecular events driving mammalian development. Stereo-seq is a spatially resolved transcriptomic technology with high resolution, sensitivity and large field of view. These methodological parameters are all fundamental for reliably profiling the transcriptomic heterogeneity of complex tissues and organisms such as mammalian embryos. Whole mouse embryo-scale spatially resolved transcriptomes have been reported recently (Liu et al., 2020; Srivatsan et al., 2021) but they contained low density of detected signals and were restricted to only one stage of organogenesis. The sparsity of gene detection in those studies makes it difficult to analyze gene expression gradients across the developing tissues and, thus, it is hard to recapitulate fine anatomical structures. We have used Stereo-seq to create the first version of MOSTA (https://db.cngb.org/stomics/mosta/), an expandable panoramic high-definition transcriptomic resource of mouse organogenesis. MOSTA provides detailed topographic information about the stepwise formation of tissues and the emergence of tissue-specific cell identities across different stages of mouse organogenesis. Examples of the latter include the identification of incipient areas of neuroblast specification in E9.5 brain subregions and of gradients of neuronal specification in the neocortex. One caveat of MOSTA is that some tissues such as the thyroid are underrepresented due to the sectioning procedure, whereas others will benefit from additional sections to fully capture the intricacy of developmental processes. It will also be important to include whole neonatal and adult mice for additional comparisons. In the future, we will add these samples and expand the number of sections with the goal of ultimately achieving 3D-transcriptomic reconstructions. Facilitating this goal, the precise spatial coordinates retrieved by Stereo-seq allow optimal integration of different slides, which might not be achieved with accuracy by techniques using randomly distributed beads. Moreover, being based on standard DNB technology it is relatively easy to enhance the sample throughput using Stereo-seq and systematically profile hundreds of tissue slices in a relatively short time.

To demonstrate the utility of MOSTA for inferring the molecular logic behind developmental processes, we have for the first time systematically dissected the spatially resolved gene regulatory networks driving organogenesis. This has identified tissue- and area-specific regulons across different embryonic stages and focusing on the brain we have also shown the correspondence with signaling cues responsible for inducing their diversification. In addition to understanding development, knowledge of these regulons and morphogens might be useful to devise or improve ways to reprogram somatic cells into other lineages *in vivo* or *in vitro* and for fine-tuning differentiation protocols using pluripotent stem cells (Wang et al., 2021). We have also projected the expression of developmental disease loci (including among others monogenic diseases involving transcription factors or morphogens and complex traits like autism) onto our spatial maps, highlighting unexpected associations such as the localization of autism-related genes in the muscle or the liver. Although mice and humans differ in multiple aspects of normal physiology or disease and caution should be taken with the interpretations, this approach has a tremendous potential to uncover novel disease mechanisms. Systematic independent explorations using our interactive portal will provide users worldwide with a unique chance to unlock the black boxes of mouse embryogenesis and developmental diseases. In this regard, expansion with datasets from genetically engineered mice that mimic developmental diseases or bear alteration of developmental pathways will be useful to test or refine any generated models of normal and abnormal development generated with MOSTA.

Importantly, larger Stereo-seq chips could enable the profiling of embryos from other mammalian species including primates. Forthcoming optimizations of Stereo-seq will also increase gene and transcript capture, facilitating the assignment of individual cell identities. Similarly, oligonucleotide-labeled antibodies (Stoeckius et al., 2017) could be incorporated into the DNB for simultaneous protein detection of surface markers, which would be useful for identifying specific cell types (e.g., infiltrating immune cells). Likewise, we anticipate that Stereo-seq will have other applications beyond RNA-sequencing such as spatially resolved epigenomics (e.g., chromatin accessibility and DNA methylation profiling). Accordingly, besides development, Stereo-seq and its future technical variations will be highly transformative for multiple research fields across kingdoms of life. Given its characteristics and applicability Stereo-seq also has the potential to move into the routine clinical practice as an extraordinary diagnostic tool complementary to medical imaging and histopathology data.

## ACKNOWLEDGMENTS

We thank the MOTIC China Group CO., Ltd. for providing technical support, and Rongqin Ke (Huaqiao University) and Jiazuan Ni (Shenzhen University) for providing technical advice. We thank Jonathan S. Weissman (Whitehead Institute) for providing helpful comments. This work was supported by the Shenzhen Key Laboratory of Single-Cell Omics (ZDSYS20190902093613831) and the Guangdong Provincial Key Laboratory of Genome Read and Write (2017B030301011); Longqi Liu was supported by the National Natural Science Foundation of China (No.31900466); Miguel A. Esteban’s lab at the Guangzhou Institutes of Biomedicine and Health was supported by the Strategic Priority Research Program of the Chinese Academy of Sciences (XDA16030502).

## AUTHOR CONTRIBUTIONS

J.W., X.X., A.C., L.L., and Y.Li conceived the idea; J.W., X.X., A.C., L.L., Y. Li and M.A.E. supervised the work; A.C., S. Liao., L.L. and M.C. designed the experiment; S. Liao, J.Y. and W.L. generated the Stereo-seq chip with the help from X.C., X.H.^1,3^ and D.F.; M.C., J.X., H.Z., H.X., L.H., P.G. and F.L. performed the majority of the experiments; K.M., L.W., X.L., X.T., H.L. and M. Li performed data preprocessing and quality evaluation; L.W., Y.Lai, X.Q., X. W. and S.H. analyzed the data; Z.L., Y.J., J.P., S.Liu, C.L., Zhi.Wang, Y.Yuan., Y.H., Q.L., Zhao.Wang, X.H.^1^, F.Z., H.K., O.W., B.W. and H.L. provided technical support; G.V., C.W., , M.N., W.Z., F.M., Y.Yin, H.Y., P.M.-C., Q.W., J.M., J.P.T, M.Lisby, R.J.C and M.U. gave the relevant advice; M.A.E., J.W., X.X., L.L., Y.Lai, W.L. and A.C. wrote the manuscript.

## RESOURCE AVAILABILITY

### Lead contact

Further information and requests for the resources and reagents may be directed to the corresponding author Xun Xu

### Material availability

All materials used for Stereo-seq are commercially available.

### Data and code availability

All raw data generated by Stereo-seq have been deposited to CNGB Nucleotide Sequence Archive (accession code: CNP0001543 (https://db.cngb.org/search/project/CNP0001543). All data were analyzed with standard programs and packages. Custom code using open-source software supporting the current study is available from the corresponding authors on request and will be uploaded at https://github.com/BGIResearch/handleBam.

## METHODS

### Tissue collection

All relevant procedures involving animal experiments presented in this paper are compliant with ethical regulations regarding animal research and were conducted under the approval of the Animal Care and Use committee of the Guangzhou Institutes of Biomedicine and Health, Chinese Academy of Sciences (license number IACUC2021002). Mouse olfactory bulb and brain were dissected from 12-week-old C57BL/6J female mice. E9.5, E10.5, E11.5, E12.5, E13.5, E15.5 and E16.5 embryos were collected from pregnant C57BL/6J female mice. After collection, tissues were snap-frozen in liquid nitrogen prechilled isopentane in Tissue-Tek OCT (Sakura, 4583) and transferred to a −80℃ freezer for storage before the experiment. Cryosections were cut at a thickness of approximately 10 μm in a Leika CM1950 cryostat; mouse olfactory bulb and mouse brain were cut coronally, mouse embryos were cut sagittally.

### Stereo-seq chip preparation

#### Generation of Stereo-seq chips

To generate the patterned array, we first synthesized two oligo sequences: one containing 25 random deoxynucleotides (nt) (TGTGAGCCAAGGAGTTGNAACTGCTGACGTACTGAGAGGCATGGCGACCTTATCAGNNNNNNNNNNNNNNNNNNNNNNNNNTTGTCTTCCTAAGACC G, Sangon) and the other a fixed sequence with phosphorylation (/5phos/CTTGGCCTCCGACTTAAGTCGGATCGTAGCCATGTCGTTC, Sangon). These two oligos were ligated by incubation with T4 ligase (NEB; 1U/μl T4 DNA ligase and 1× T4 DNA ligation buffer) and splint oligo (TCGGAGGCCAAGCGGTCTTAGGAA, Sangon, 1 μM) at 37℃ for 2 hours. The products were purified using the Ampure XP Beads (Vazyme, N411-03) and then PCR amplified with the following steps: 95℃ for 5 minutes, 12 cycles at 98℃ for 20 seconds, 58℃ for 20 seconds, 72℃ for 20 seconds and a final incubation at 72℃ for 5 minutes. PCR products were purified using the Ampure XP Beads (Vazyme, N411-03). DNBs were then generated by rolling circle amplification and loaded onto the patterned chips according to the MGI DNBSEQ-Tx sequencer manual. Next, to determine the distinct DNB-CID sequences at each spatial location, single-end sequencing was performed using a sequencing primer (CTGCTGACGTACTGAGAGGCATGGCGACCTTATCAG, Sangon, 1 μM) in a MGI DNBSEQ-Tx sequencer with SE25 sequencing strategy. After sequencing, the capture oligo (/5phos/TTGTCTTCCTAAGACNNNNNNNNNNTTTTTTTTTTTTTTTTTTTTTV, Sangon, 1 μM) including 22 nt poly-T and 10 nt UMI was hybridized with the DNB in 5× SSC buffer at 37℃ for 30 minutes, and then incubated with T4 ligase (NEB, 1 U/μl T4 DNA ligase, 1× T4 DNA ligation buffer and 0.5% PEG2000) at 37℃ for 1 hour. This procedure produces capture probes containing a 25 nt CID barcode, a 10 nt UMI and a 22 nt poly-T ready for poly-A RNA capture. A schematic describing this procedure is included in the MOSTA website (https://db.cngb.org/stomics/mosta/).

#### Calling of CID

CID sequences together with their corresponding coordinates for each DNB were determined using a base calling method according to manufacturer’s instruction of MGI DNBSEQ-Tx sequencer. After sequencing, the capture chip was split into smaller size chips (5 mm × 10 mm, 10 mm × 10 mm, 10 mm × 20 mm) ready for use. At this stage, we filtered out all duplicated CID that corresponded to non-adjacent spots.

### Stereo-seq library preparation and sequencing

#### Tissue processing

Tissue sections were adhered to the Stereo-seq chip surface and incubated at 37℃ for 3 minutes. Then, they were fixed in methanol and incubated for 30 minutes at −20℃, after which they were ready to be used for Stereo-seq. Where indicated, the same sections were stained with nucleic acid dye (Thermo fisher, Q10212), or adjacent sections were subjected to tissue histology examination using hematoxylin and eosin staining according to standard procedure. Imaging for both procedures was performed with a Ti-7 Nikon Eclipse microscope.

#### In situ reverse transcription

Tissue sections placed on the chip were permeabilized using 0.1% pepsin (Sigma, P7000) in 0.01 M HCl buffer, incubated at 37℃ for 12 minutes and then washed with 0.1× SSC buffer (Thermo, AM9770) supplemented with 0.05 U/μl RNase inhibitor (NEB, M0314L). RNA released from the permeabilized tissue and captured by the DNB was reverse transcribed overnight at 42℃ using SuperScript II (Invitrogen, 18064-014, 10 U/μl reverse transcriptase, 1 mM dNTPs, 1 M betaine solution PCR reagent, 7.5 mM MgCl_2_, 5 mM DTT, 2 U/ml RNase inhibitor, 2.5 μM Stereo-TSO and 1× First-Strand buffer). After reverse transcription, tissue sections were washed twice with 0.1× SSC buffer and digested with Tissue Removal buffer (10 mM Tris-HCl, 25 mM EDTA, 100 mM NaCl and 0.5% SDS) at 37℃ for 30 minutes. cDNA-containing chips were then subjected to Exonuclease I (NEB, M0293L) treatment for 1 hour at 37℃ and were finally washed once with 0.1x SSC buffer.

#### Amplification

The resulting cDNA were amplified with KAPA HiFi Hotstart Ready Mix (Roche, KK2602) with 0.8 μM cDNA-PCR primer. PCR reactions were conducted as follows: first incubation at 95℃ for 5 minutes, 15 cycles at 98℃ for 20 seconds, 58℃ for 20 seconds, 72℃ for 3 minutes and a final incubation at 72℃ for 5 minutes.

#### Library construction and sequencing

The concentrations of the resulting PCR products were quantified by Qubit™ dsDNA Assay Kit (Thermo, Q32854). A total of 20 ng of DNA were then fragmented with *in-house* Tn5 transposase at 55℃ for 10 minutes, after which the reactions were stopped by the addition of 0.02% SDS and gently mixing at 37 ℃ for 5 minutes after fragmentation. Fragmentation products were amplified as described below: 25 ml of fragmentation product, 1× KAPA HiFi Hotstart Ready Mix and 0.3 μM Stereo-Library-F primer, 0.3 μM Stereo-Library-R primer in a total volume of 100 ml with the addition of nuclease-free H_2_O. The reaction was then run as: 1 cycle of 95℃ 5 minutes, 13 cycles of (98℃ 20 seconds, 58℃ 20 seconds and 72 ℃ 30 seconds) and 1 cycle of 72 ℃ 5 minutes. PCR products were purified using the Ampure XP Beads (Vazyme, N411-03) (0.6× and 0.15×), used for DNB generation and finally sequenced (35 bp for read 1, 100 bp for read 2) on a MGI DNBSEQ-Tx sequencer.

### Stereo-seq raw data processing

Fastq files were generated using a MGI DNBSEQ-Tx sequencer. CID and UMI are contained in the read 1 (CID: 1-25bp, UMI: 26-35bp) while the read 2 consist of the cDNA sequences. CID sequences on the first reads were first mapped to the designed coordinates of the *in situ* captured chip achieved from the first round of sequencing, allowing 1 base mismatch to correct for sequencing and PCR errors. Reads with UMI containing either N bases or more than 2 bases with quality score lower than 10 were filtered out. CID and UMI associated with each read were appended to each read header. Retained reads were then aligned to the reference genome (mm10) using STAR (Dobin et al., 2013) and mapped reads with MAPQ ≥10 were counted and annotated to their corresponding genes using *handleBam* (available at https://github.com/BGIResearch/handleBam). UMI with the same CID and the same gene locus were collapsed, allowing 1 mismatch to correct for sequencing and PCR errors. Finally, this information was used to generate a CID-containing expression profile matrix.

### Unsupervised spatially-constrained clustering of Stereo-seq data

Expression profile matrix was divided into non-overlapping bins covering an area of X × X DNB, with X ∈ (*1, 3, 14, 50, 140*) and the transcripts of the same gene aggregated within each bin. After this step, data were log-normalized in Scanpy and spatially-constrained unsupervised clustering was performed. In brief, to include spatial information during clustering, we first built a spatial k-nearest neighbor graph 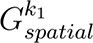 (*k*_1_ is by default set to be 8 as each bin has 8 nearest spatial neighbors) using Squidpy and then took the union with the k-nearest neighbor graph 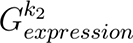 based on transcriptomic data ( 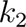 is by default set to be 30). The combined graph ( 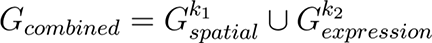) was then used as input for leiden clustering. Further, each cluster was annotated based on the cluster specific markers identified by the rank_genes_groups function of scanpy using default parameters as well as the anatomical annotation based on eHistology Kaufman Annotations (http://www.emouseatlas.org/emap/home.html) or Allen Brain Atlas (http://mouse.brain-map.org/).

### Comparison of Stereo-seq with other published methods

HDST data (Vickovic et al., 2019) were taken from GSE130682, SLIDE-seqV2 data (Stickels et al., 2021) from the Single Cell Portal of the Broad Institute, DBiT-seq data (Yao et al., 2020) from GSE137986, and Visium data (Lebrigand et al., 2020b) from GSE153859, Seq-Scope data were taken from GSE169706 (Cho et al., 2021a). For Fig. 1c, to ensure that proper comparisons were made, the data of Stereo-seq were binned into bin 3 (3 × 3 resolution, ∼2 μm diamater), bin 14 (14 × 14 resolution, ∼10 μm diamater) or bin 140 (140 × 140 resolution, ∼100 μm diamater). For **Extended Data Fig. 1e**, data from HDST were binned into bin 5 (5 × 5, ∼10 μm diamater).

### Spatially-resolved gene regulatory networks

The analysis of regulon activity was performed by following the standard SCENIC pipeline (Aibar et al., 2017). The input to SCENIC was the bin 50 (50 x 50 resolution) expression matrix. The expression matrix was loaded into GENIE3 and the co-expressed gene network for each transcription factor was constructed. Transcription factor co-expression modules were then analyzed by RcisTarget and their potential targets further filtered by default parameters. Filtered potential targets were used to build the regulons. Regulon activity (Area Under the Curve) was analyzed by AUCell and the active regulons were determined by AUCell default threshold. The activity of regulons for each bin was then mapped to the physical space. The gene network for Figure 3F was constructed by selecting the target genes of corresponding transcription factors related to neuronal development, and further visualized by Cytoscape.

### Identification of spatially auto-correlated gene or regulon modules

Spatially auto-correlated gene or regulon modules were identified using Hotspot (DeTomaso and Yosef, 2021). The expression matrix for the top 5,000 variable genes and all regulon activity matrix of each embryo were used as input. For gene module, the data were first normalization by the total UMI number of each bin, then create knn graph of genes using the create_knn_graph function with the parametes: n_neighbors = 30 (for regulon, n_neighbors = 10), then genes or regulons with significant spatial autocorrelation (FDR < 0.05) were kept for further analysis. And the modules were identified using the create_modules with the parameters: min_gene_threshold = 20 and fdr_threshold = 0.05 (for the regulon: min_gene_threshold = 5 and fdr_threshold = 0.05).

### Deconvolution of cell types

We applied SPOTlight (Elosua-Bayes et al., 2021) to deconvolute and map all cell types in the murine telencephalon. Mouse embryonic cortex scRNA-seq data were retrieved from GSE153164 (Di Bella et al., 2021). We chose cell type with the highest probability out of all cell types provided by the spotlight_deconvolution function with commendatory parameters as the final cell type for each bin. The cell type specific proportions were projected to the physical space.

### Rostral-caudal brain gene expression dynamics

The CP of E15.5 telencephalon was manually dissected into 16 blocks along the rostral-caudal direction based on spatial coordinates. Variable genes along the rostral to caudal axis were identified using the differentialGeneTest function of monocle (Qiu et al., 2017) based on the corrdinates of the blocks. Then DEG along the rostral-caudal axis were identified with *p* < 0.05.

### RNA velocity analysis

Analysis was performed using Dynamo (Qiu et al., 2021) following the tutorial found at https://dynamo-release.readthedocs.io/. Unspliced and spliced RNA for each bin (20 × 20 resolution) were extracted from E15.5 telencephalon region with handleBam. The data matrix was then processed by Dynamo to normalize the expression, select feature genes and perform dimension reduction via UMAP, followed by default parameters to estimate the kinetic parameters and gene-wise RNA velocity vectors that were then projected to the physical space. Specifically, the new “Fokker-Planck” kernel implemented in Dynamo instead of the conventional correlation or cosine kernel was used for the projection of high-dimensional RNA velocity vectors to physical space. Streamlines were used to visualize the velocity vector flows on telencephalon in which only velocity flows for the relevant domains are visualized. To facilitate the understanding of gene expression dynamics over space, we first reconstructed the continuous vector field in the UMAP space, and then utilized a new method, hodge decomposition, implemented in Dynamo that takes the simplicial complexes (a sparse directional graph) constructed based on the learned vector field function to infer the vector field based pseudotime (vf pseudotime). At last, we also illustrated the kinetics of all genes with highest PCA loading as well as the DEG identified by spatially-constrained clustering along the vf pseudotime.

### Spatial gene enrichment analysis of human developmental disease associated genes

A list of 1,940 genes corresponding to monogenic diseases was retrieved from DDG2P (version 2.29). The gene set related to autism were taken from MSigDB (Liberzon et al., 2015). For monogenic disease analysis, genes with depth-normalized expression value greater than 1(729 genes) were kept for further analysis. The expression level of indicated organs at selected time point were aggregated, Z score normalized, and further visualized by heatmap.

---

**Figure S1.**
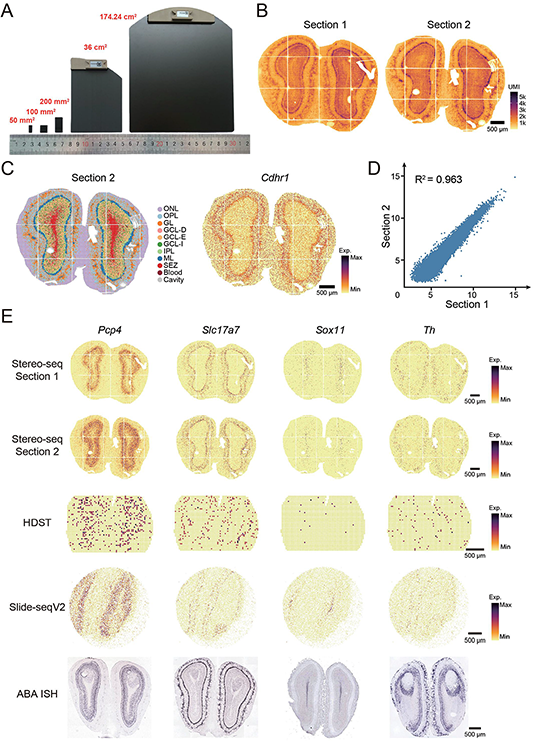
Size of stereo-seq chips and assessment of performance, related to Figure 1. **A.** Stereo-seq chips of different sizes ranging from 50 mm^2^ to 174.24 cm^2^. **B.** Spatial visualization of the number of UMI captured by Stereo-seq from two adjacent mouse olfactory bulb coronal sections at bin 14 resolution. **C.** Unsupervised spatially-constrained clustering of the mouse olfactory bulb section (section 2) at bin 14 resolution. **D.** Pearson correlation coefficient (R^2^ = 0.9666) of pseudo-bulk profiles from the two Stereo-seq experiments of mouse olfactory bulb. **E.** Spatial visualization of the indicated genes in the two independent Stereo-seq experiments for mouse olfactory bulb and reported HDST, Slide-seqV2 and ISH datasets (Lein et al., 2007; Stickels et al., 2021; Vickovic et al., 2019). Scale bars, 500 μm.

**Figure S2.**
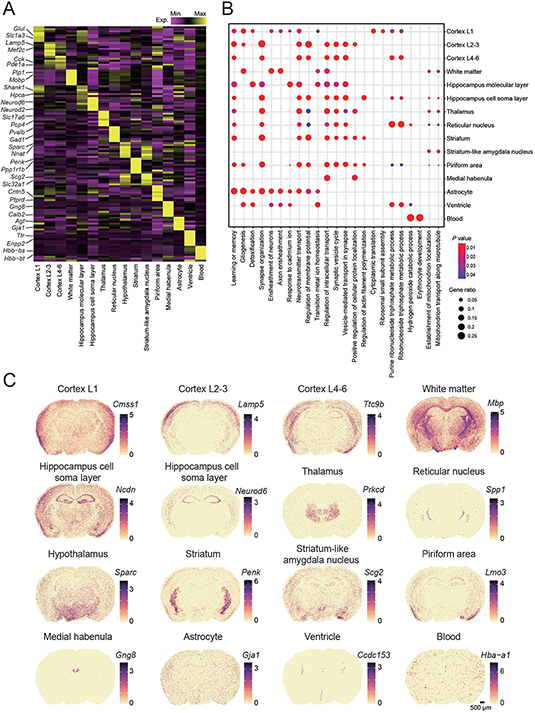
Stereo-seq detects different anatomical structures in the adult mouse brain, related to Figure 1. **A.** Heatmap showing the mean expression level of DEG between the indicated anatomical structures identified in Figure 1F. **B.** GO analysis of DEG between the indicated anatomical structures. Selected GO terms (Benjamini-Hochberg corrected *P* value) are shown. **C.** Spatial visualization of the expression of indicated genes representing different anatomical structures in a whole adult mouse brain coronal section analyzed by Stereo-seq at bin 50 resolution. Gene expression levels were quantified as log_2_ (transcripts per 10K UMI + 1), which applies to all similar plots.

**Figure S3.**
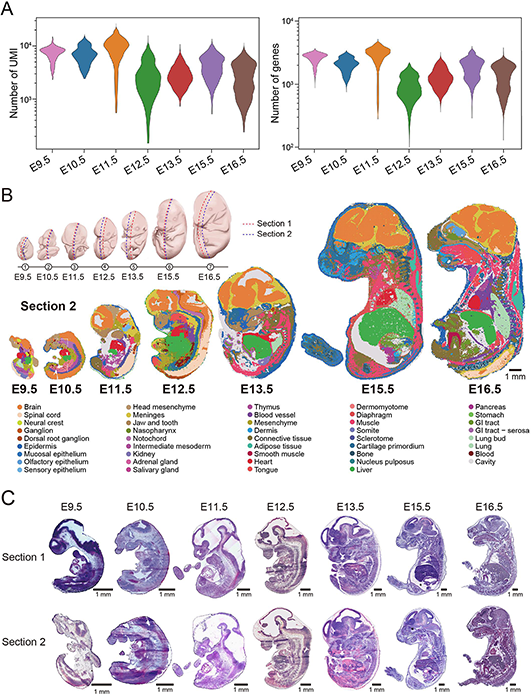
Spatially resolved transcriptomic atlas of mouse organogenesis, related to Figure 2. **A.** Violin plot showing the number of transcripts (left) and genes (right) captured by Stereo-seq at different stages of embryonic development, related to Figure 2A. **B.** Top: overview of the sampled embryonic time points and sections. Middle and bottom: spatially-constrained unsupervised clustering of additional embryo sections across E9.5, E10.5, E11.5, E12.5, E13.5, E15.5, and E16.5 also identifies anatomical structures (annotation is shown below). Bins are colored according to different anatomical structures. Scale bar, 1 mm. **C.** Hematoxylin and eosin staining of embryo sections adjacent to the corresponding Stereo-seq sections. Scale bars, 1 mm.

**Figure S4.**
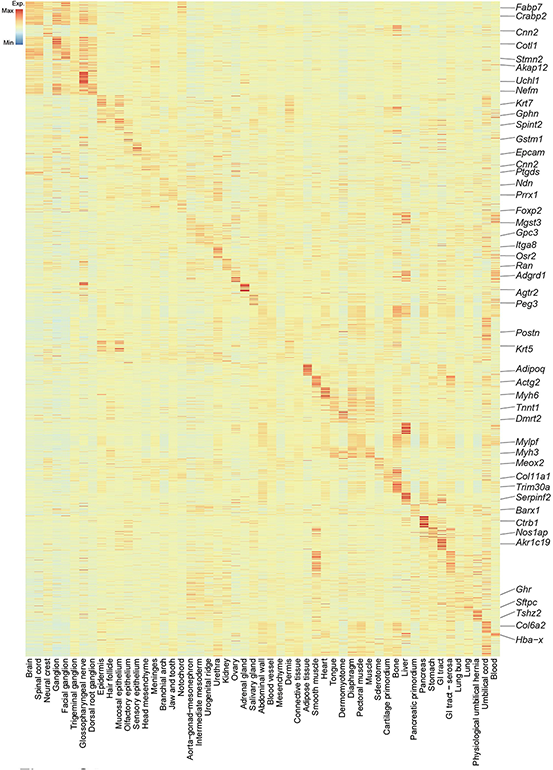
DEG of mouse embryo anatomical structures, related to Figure 2. Heatmap showing the normalized expression of DEGs for the indicated anatomical structures of the mouse embryo sections profiled by Stereo-seq in Figure 2A.

**Figure S5.**
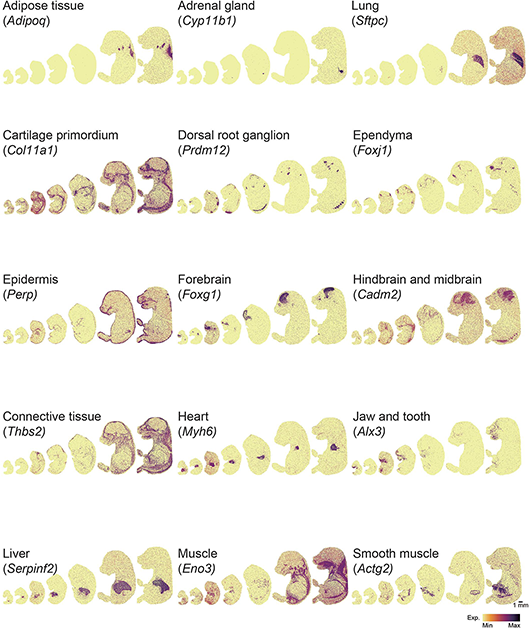
Expression of selected markers for specific mouse embryo tissues, related to Figure 2. Spatial visualization of the expression of DEGs enriched in the indicated tissues of the mouse embryo sections profiled by Stereo-seq in Figure 2A.

**Figure S6.**
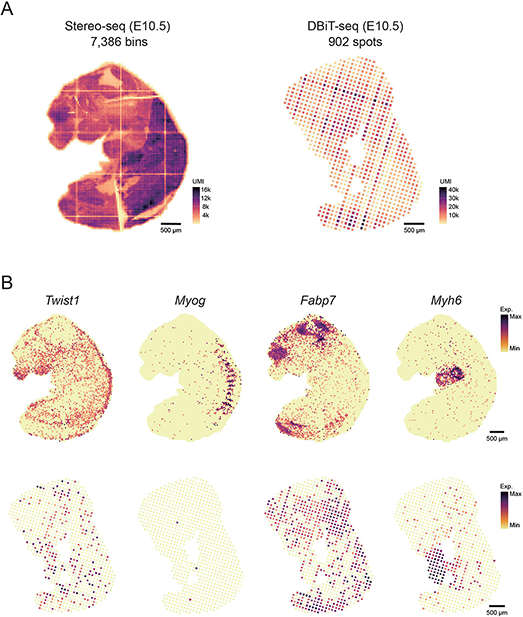
Reanalysis of a mouse E10 embryo dataset generated by DBiT-seq, related to Figure 2. **A.** Spatial heatmap indicating the number of UMI captured by Stereo-seq from an E10.5 embryo section (same as in Figure 2A) at bin 50 (left) compared to an E10 DBiT-seq dataset at the same resolution (right). Scale bar, 500 μm. **B.** Spatial visualization of the expression of the indicated genes in Stereo-seq (upper panels) and DBiT-seq (lower panels) mouse embryo datasets. Scale bars, 500 μm.

**Figure S7-S12.**
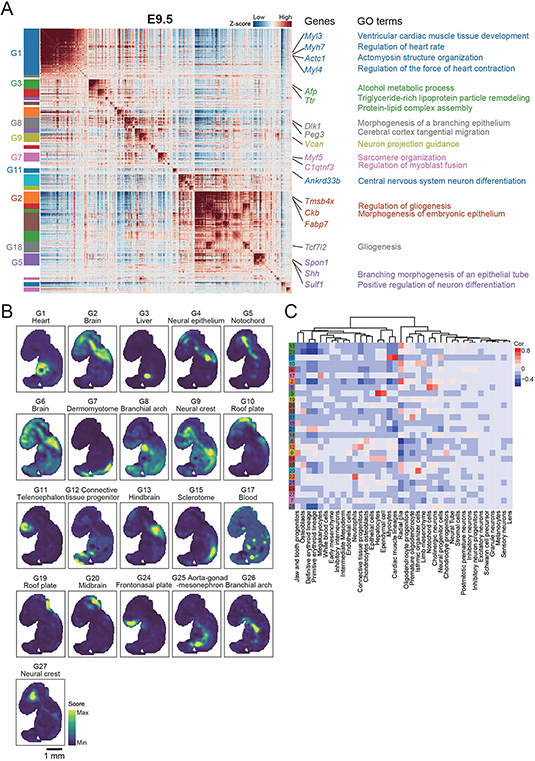
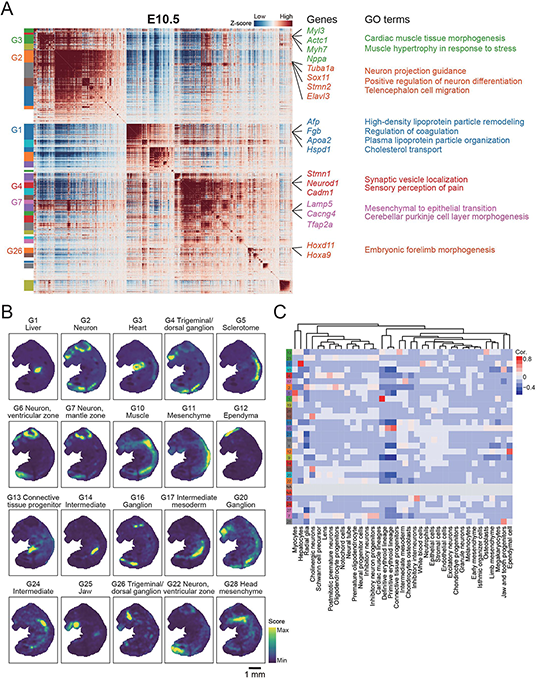
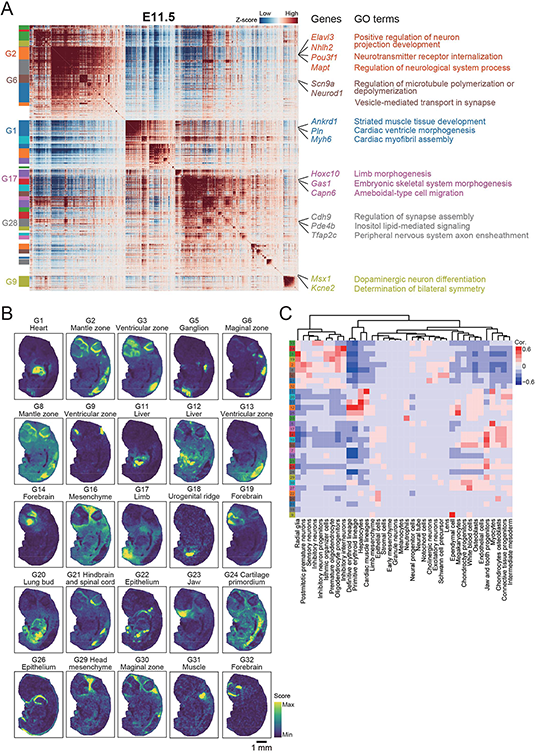
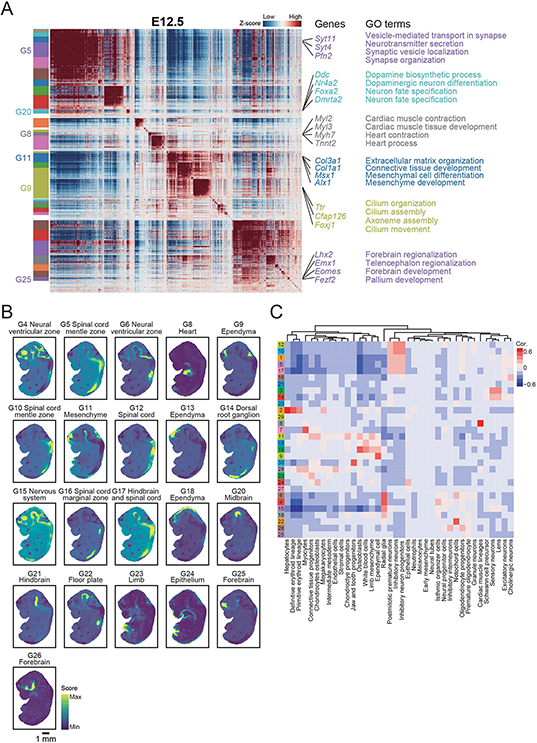
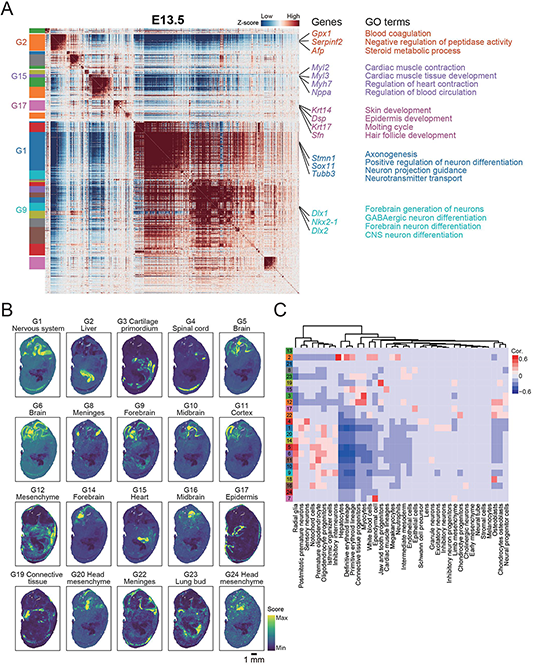
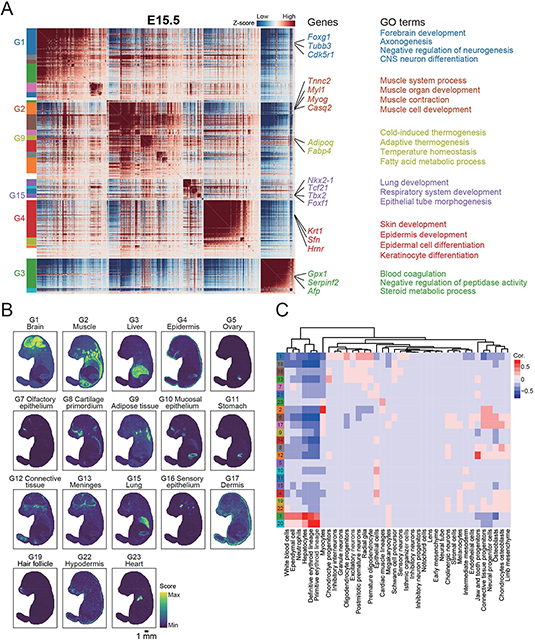
Hotspot identified spatial gene expression modules in E9.5-E15.5 mouse embryo sections, related to Figure 2. **A.** Heatmap showing the genes with significant spatial autocorrelation (2,082, 1,916, 3,062, 2,831, 2,074, 2,480 genes, FDR < 0.05) grouped into 28, 28, 33, 30, 24 and 23 gene modules based on pairwise spatial correlations of gene expression in the same embryo sections of Figure 2A from E9.5 to E15.5, respectively. Selected genes and GO terms related to representative modules are highlighted on the right side. **B.** Heatmap showing the Pearson correlation of module score for each spatial autocorrelation module and the expression of sets of specific genes for each cell type from a reported dataset (Cao et al., 2019). **C.** Spatial visualization of gene modules related to the indicated anatomical structures. Scale bar, 1 mm.

**Figure S13.**
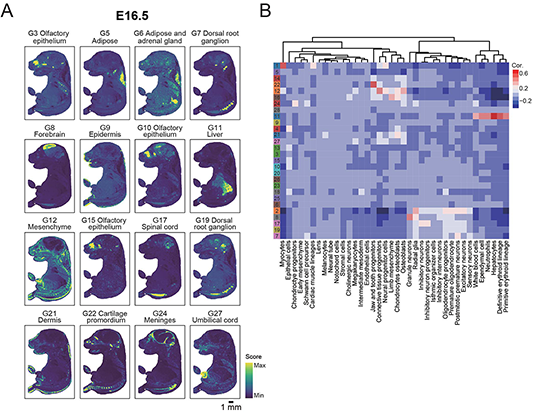
Hotspot identified spatial gene expression modules in an E16.5 mouse embryo sections, related to Figure 2. **A.** Heatmap showing the Pearson correlation of module score for each spatial autocorrelation module and the expression of sets of specific genes for each cell type from a reported dataset (Cao et al., 2019). **B.** Spatial visualization of gene modules related to the indicated anatomical structures. Scale bar, 1 mm.

**Figure S14-S19.**
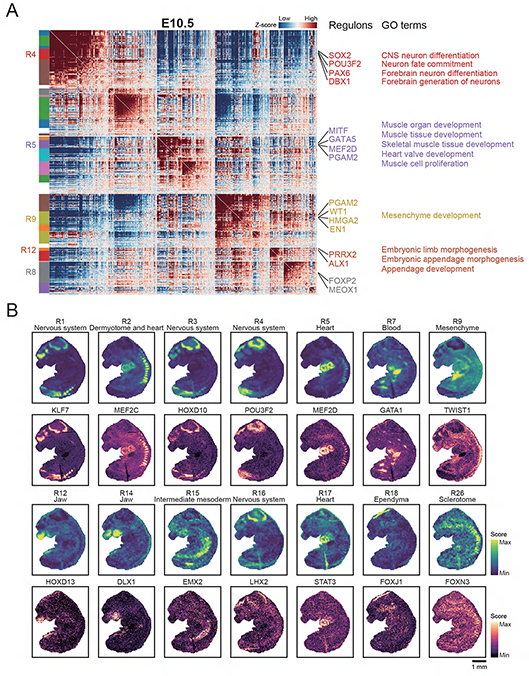
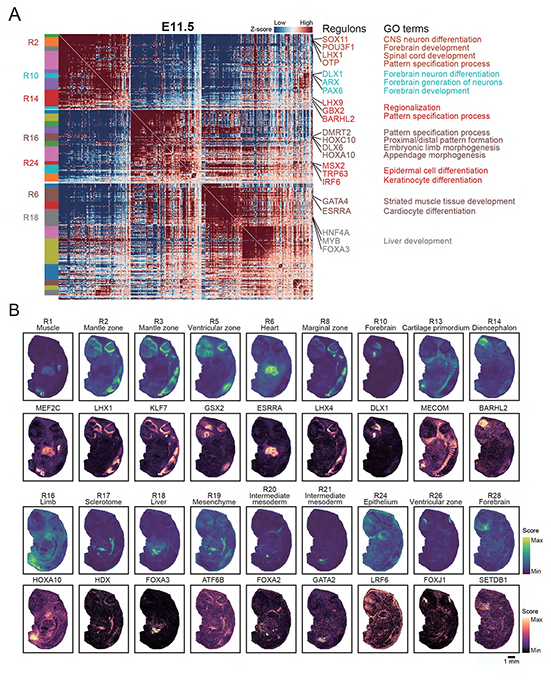
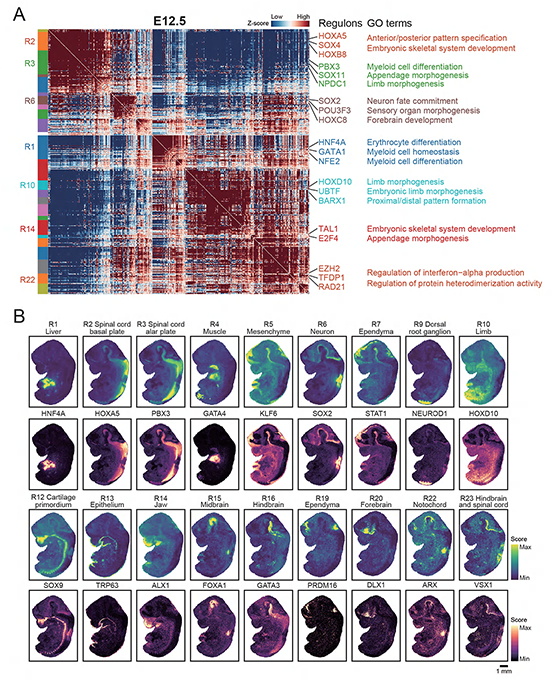
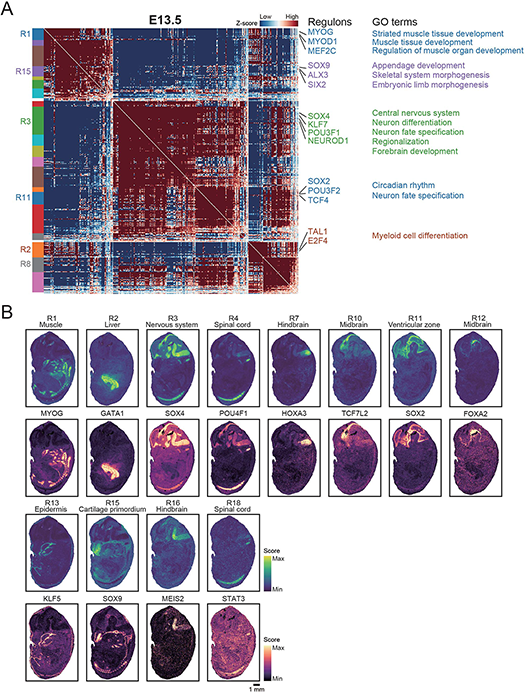
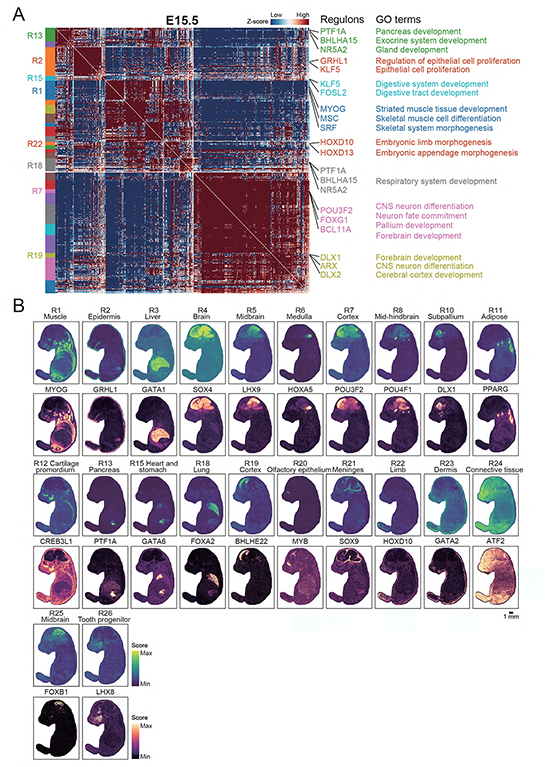
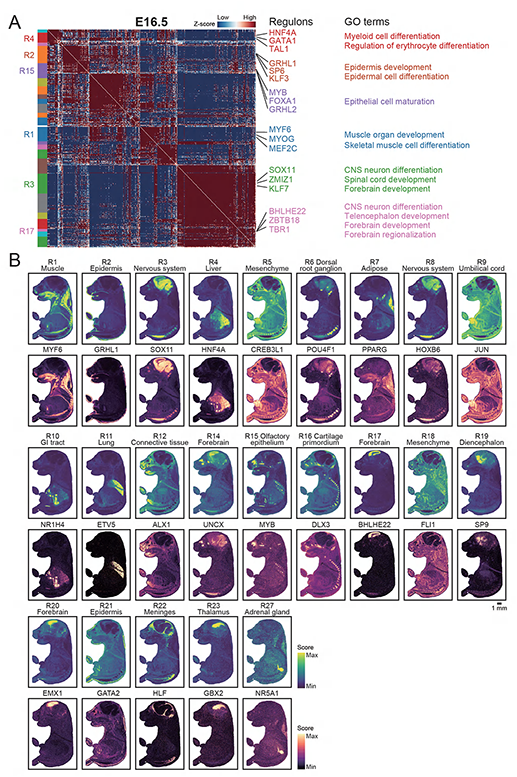
Hotspot identified spatial regulatory modules in E10.5-E16.5 mouse embryo sections, related to Figure 3. **A.** Heatmap showing the regulons with significant spatial autocorrelation (553, 554, 500, 380, 467 and 480 regulons, FDR < 0.05) grouped into 26, 31, 25, 20, 28 and 27 regulon modules based on pairwise spatial correlations of regulon activity in the same embryo sections of Figure 2A from E10.5 to E16.5, respectively. Selected regulons related to representative modules and their corresponding GO terms are shown. **B.** Spatial visualization of the indicated regulon modules and their representative regulon for different anatomical structures. Scale bar, 1 mm.

**Figure S20.**
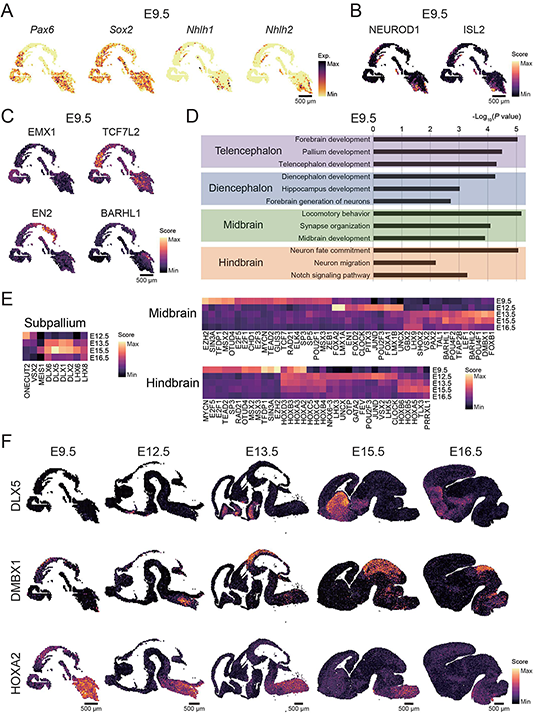
Stereo-seq reconstructs the spatiotemporal gene regulatory network of the developing mouse brain, related to Figure 3. **A.** Spatial visualization of the expression of the indicated genes representing neuroepithelium (left) and early neuroblasts in the E9.5 (right) brain. **B.** Spatial visualization of the indicated regulon activity in early neuroblasts of the E9.5 brain. **C.** Spatial visualization of the indicated regulon activity representing different anatomical structure of the E9.5 brain. **D.** GO analysis of target genes for transcription factors corresponding to different anatomical structures of the E9.5 brain as in Figure 3F. **E.** Heatmap showing the normalized regulon activity of top transcription factors corresponding to the subpallium, midbrain, and hindbrain development of mouse brain. **F.** Spatial visualization of the indicated regulon activity for the subpallium (DLX5), midbrain (DMBX1) and hindbrain (HOXA2). Scale bars, 500 μm.

**Figure S21.**
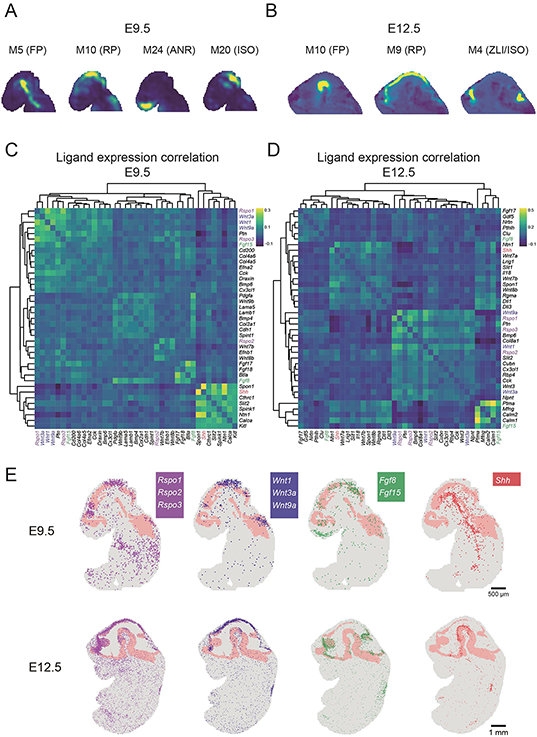
Characterization of morphogens expressed by secondary organizer centers in the developing mouse brain, related to Figure 3. **A** and **B**. Spatial visualization of the indicated modules representing secondary organizer centers of the E9.5 and E12.5 brain. **C** and **D**. Heatmap showing the Spearman’s correlation of ligands enriched in the secondary organizes centers of the E9.5 and E12.5 brain. **E**. The spatial scatter plots showing single DNB that captured the transcripts of the indicated morphogens at the ISO, ANR, ZLI, RP, and FP organizer centers at E9.5 (bottom left) and E12.5 (bottom right) embryos. Plots were superimposed with E9.5 or E12.5 embryo structures. Scale bar, 500 μm.

**Figure S22.**
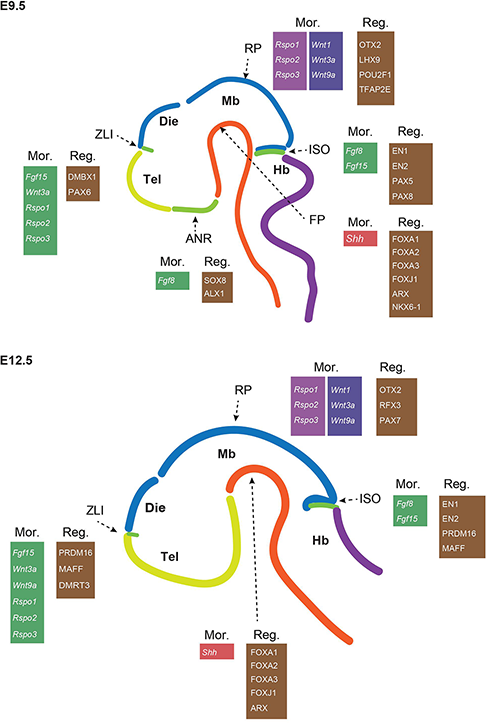
Schematic representation of morphogens and regulons at the indicated organizer centers, related to Figure 3.

**Figure S23.**
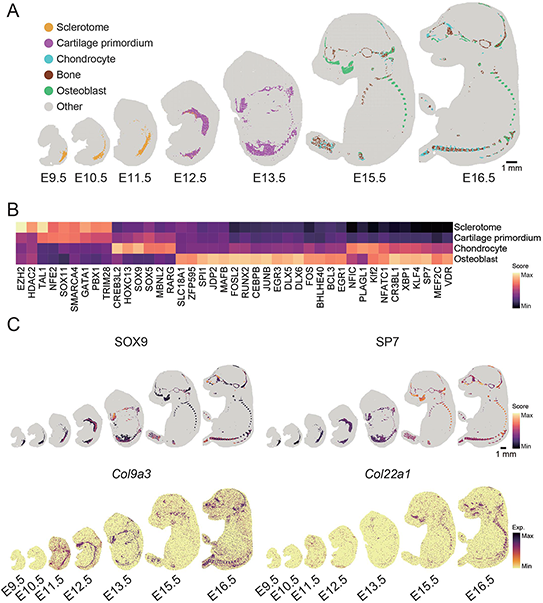
Stereo-seq reconstructs the spatiotemporal gene regulatory networks of the developing mouse skeleton, related to Figure 3. **A.** Unsupervised spatially-constrained clustering of the developing mouse skeleton based on regulon activity. **B.** Heatmap showing the normalized regulon activity score of the top transcription factors enriched in the indicated anatomical structures during mouse embryonic skeleton development. **C.** Spatial visualization of the activity of the indicated regulons for chondrocytes (SOX9) and osteoblasts (SP7). **D.** Spatial visualization of the expression of the indicated genes representing the early (*Col9a3*) and late (*Col22a1*) stages of ossification. Scale bar, 500 μm.

**Figure S24.**
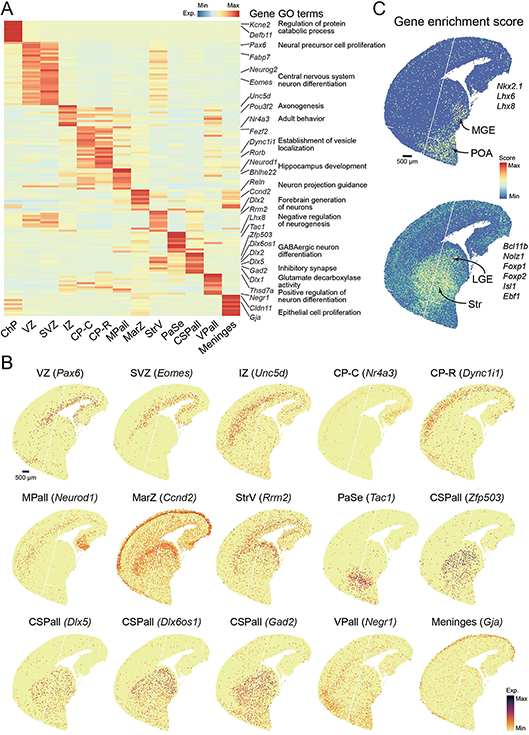
Stereo-seq reconstructs the developing mouse forebrain, related to Figure 4. **A.** Heatmap showing the normalized expression of DEGs for the indicated anatomical structures of the E15.5 forebrain in Figure 4A. **B.** Spatial visualization of indicated genes representing different anatomical structures of the E15.5 forebrain. Scale bar, 500 μm. **C.** Spatial visualization of gene enrichment score of MGE, POA, LGE and Str. Genes related to each structure are labeled on the right side. MGE, medial ganglionic eminence; POA, preoptic area; LGE, lateral ganglionic eminence; Str, striatum. Scale bar, 500 μm.

**Figure S25.**
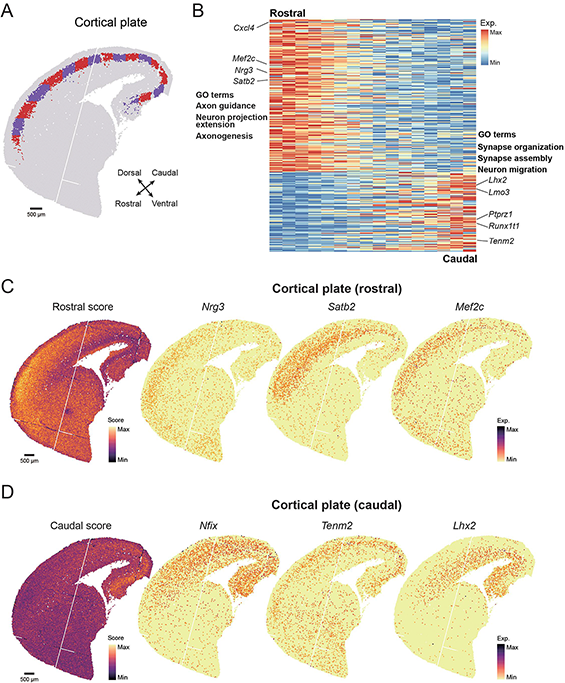
Rostral-caudal patterning of the CP, related to Figure 4. **A.** Manual dissection of the CP based on spatial coordinates. Scale bar, 500 μm. **B.** Heatmap showing the normalized expression of genes with rostral-caudal gradient. **C.** Spatial visualization of module score (left) representing the aggregated expression level of CP (rostral) specific gene set and representative genes (right). Scale bar, 500 μm. **D.** Spatial visualization of module score (left) representing the aggregated expression level of CP (caudal) specific gene set and representative genes (right). Scale bar, 500 μm.

**Figure S26.**
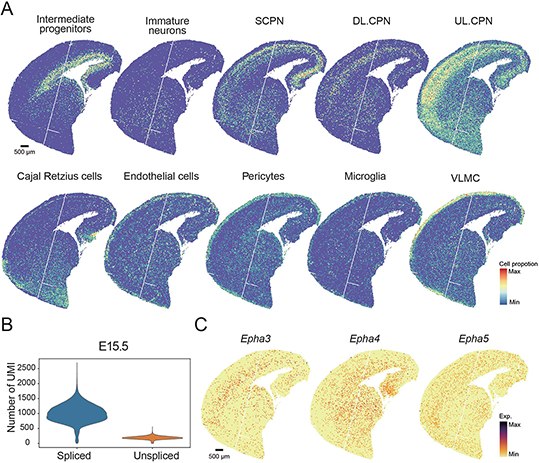
Mapping cell-type localization across forebrain, related to Figure 4. **A.** Spatial visualization of cell proportion score representing different cell types. Scale bar, 500 μm. **B.** Violin plot showing the number of spliced and unspliced transcripts. **C.** Spatial visualization of the expression of indicated genes related to tangential migration of interneurons. Scale bar, 500 μm.

**Figure S27.**
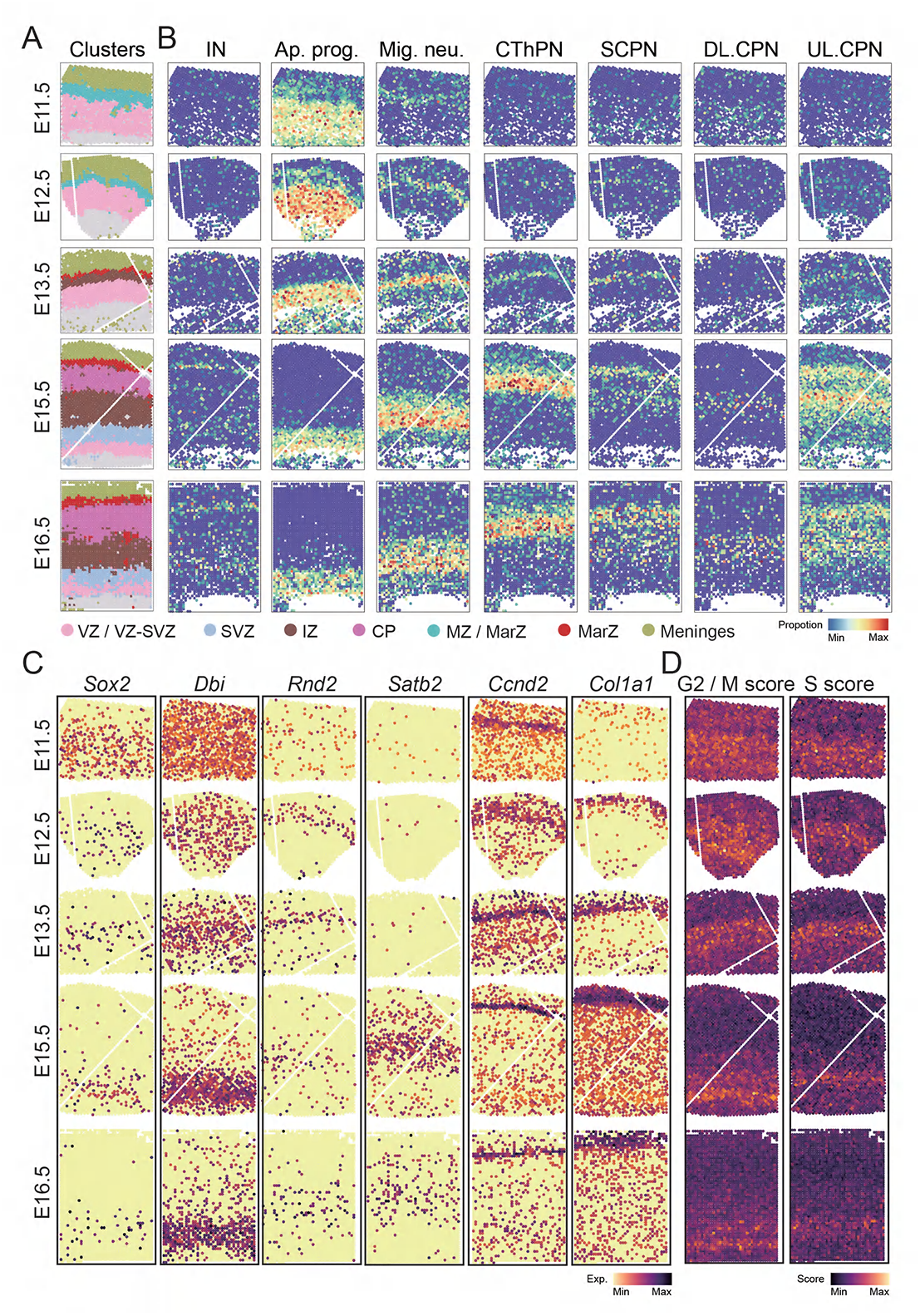
Mapping cell-type localization across neocortex development, related to Figure 4. **A.** Unsupervised spatially-constrained clustering of the cortex from E11.5, E12.5, E13.5, 15.5, and E16.5 of mouse embryo. **B.** Spatial visualization of the cell proportion score across different stages of the mouse neocortex. **C.** Spatial visualization of representative genes for VZ, SVZ, IZ, CP, MarZ and meninges across different stages of the mouse neocortex. **D.** Spatial visualization of the cell cycle score across different stages of the mouse neocortex.

**Figure S28.**
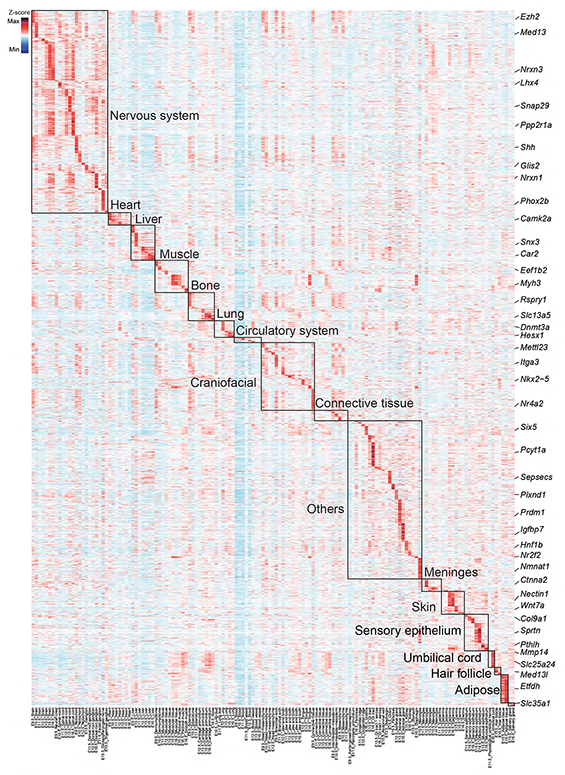
Spatial mapping of genes associated with human developmental disorders in different mouse embryo stages, related to Figure 5. Heatmap showing the normalized expression level of 1,940 genes selected from the Deciphering Developmental Disorders database in the indicated organs of E9.5 to E16.5 embryos.

**Table S1. Gene modules for each embryonic stage, related to** Figure 2.

**Table S2. Regulon modules for each embryonic stage, related to** Figure 3.

**Table S3. Expression of genes related to the rostral-caudal gradient in the cortical plate, and genes related to the pseudotime of neuronal migration, related to** Figure 4.

**Table S4. Expression of genes related to monogenic diseases and autism, related to** Figure 5.

